# Parathyroid Hormone Receptor 1 Signaling Encodes Multifaceted Metabolic Programs in AgRP Neurons

**DOI:** 10.64898/2026.10.08.757566

**Authors:** Mathieu Galerne, Eleni Siopi, Audrey Rousseaud, Léa Vinel, Nour Mesto, François Mailliet, Fabienne Liénard, Valérie Boitez, Emmanuelle Nedelec, Stéphanie Moriceau, Manon Rivagorda, Maya Faour, Julien Castel, Soham Saha, Christiane Schreiweiss, Ivan Nemazanyy, David Romeo-Guitart, Nicolas Dupont, Cyril Confavreux, Alexandre Benani, Serge Luquet, Franck Oury

**Author notes:** To whom correspondence should be addressed: Drs. Eleni Siopi and Franck Oury, INSERM U1151, Institut Necker Enfants-Malades, Team “Integrative Neurobiology” 160 rue Vaugirard, Paris Cedex 15. and. Equal contribution as co-first author (Mathieu Galerne, Eleni Siopi, Audrey Rousseaud). Equal contribution as co-second author (Léa Vinel, Nour Mesto, François Mailliet).

## Abstract

Organisms continuously adapt food intake and nutrient utilization to fluctuating energy demands, requiring the brain to accurately integrate signals coming from diverse metabolic organs and local signals into distinct physiological responses. How hypothalamic neurons discriminate these signals to independently regulate feeding behavior and peripheral metabolism remains poorly understood. In recent years, a coordinated regulation between bone physiology and the central control of energy metabolism has emerged, opening questions about potential common hormonal networks and signaling pathways between the bone and the brain. Here, we identify parathyroid hormone receptor 1 (PTH1R) signaling, a master regulator of calcium ion biochemistry and bone formation, in Agouti-related peptide (AgRP) neurons as an unexpected neuroendocrine mechanism for the control of feeding behaviourand nutrient partitioning. the two cognate endocrine and paracrine ligands, PTH and PTH-related peptide (PTHrP), selectively activat segregated intracellular kinases activation and lipophagic processes in AgRP neuron, ultimately producing convergent physiological outputs. Together, our findings identify the parathyroid hormonal system acts as a fine-tuning mechanism in specialized neuronal circuits allowing for hypothalamic adaptive responses to energy demands and systemic metabolic signals.

## Introduction

Maintaining energy homeostasis requires the brain to continuously integrate endocrine, nutrient-derived, and locally produced signals and translate this complex information into distinct physiological responses that match the organism’s metabolic demands (Dietrich & Horvath, 2013; Schwartz et al., 2000; Schwartz & Porte, 2005; Waterson & Horvath, 2015). How hypothalamic neurons decode these diverse cues to independently regulate feeding behavior, energy expenditure, nutrient utilization, and adaptive responses to fasting remains a fundamental question in neuroendocrinology.

Within the hypothalamus, the arcuate nucleus (ARC) serves as a primary integrative center for whole-body energy homeostasis. Orexigenic Agouti-related peptide (AgRP) neurons and anorexigenic pro-opiomelanocortin (POMC) neurons form the core of the melanocortin system and represent a primary substrate for the integration of a wide range of endocrine, nutritional, and locally produced signals that collectively regulate energy balance (Barsh & Schwartz, 2002; Dietrich & Horvath, 2013; Elmquist et al., 1999; Timper & Brüning, 2017). Beyond their established role in promoting feeding, accumulating evidence shows that AgRP neurons can exert a complementary yet segregated control of metabolism, including substrate utilization, fat oxidation, and fasting adaptation, indicating that these neurons can orchestrate distinct physiological outputs beyond food intake alone (Bouyakdan et al., 2022; Manceau et al., 2020). While parallel circuits organization of these neurons, together with molecularly defined sub populations of ARC neurons with specific G-protein-coupled receptor (GPCR) equipment has been proposed to contribute to this functional diversity (Betley et al., 2013), the molecular mechanisms allowing AgRP neurons to discriminate endocrine signals and engage distinct metabolic programs remain largely unknown.

Increasing evidence indicates that bone is not only a structural organ but also as an unsuspected endocrine regulator of whole-body metabolism (Ducy et al., 2000; Friedman et al, 2016; Fulzele et al., 2010; Kajimura et al., 2013; Li et al., 2019; Mosialou et al., 2017; Takahashi & Cone, 2005; Takeda et al., 2002). Bidirectional communication between the skeleton and the central nervous system coordinates bone physiology, energy balance, and glucose metabolism, suggesting that both organs share common hormonal signaling pathways (Karsenty, 2006; Karsenty & Oury, 2012; Yadav et al., 2009).

Among the principal endocrine regulators of bone physiology are parathyroid hormone (PTH) and parathyroid hormone-related peptide (PTHrP), which coordinate calcium homeostasis, bone remodeling, and skeletal development. Intriguingly, alterations in circulating PTH levels are consistently associated with obesity, insulin resistance, and type 2 diabetes (Bannon et al., 1988; Corbetta et al., 2018; Dent et al., 1962; Legakis & Mantouridis, 2009; Ljunghall et al., 1983; Martínez-García et al., 2012; Ministrini et al., 2020; Roca-Rodríguez et al., 2015; Taylor & Khaleeli, 1997; Taylor & Khaleeli, 2001; Walsh et al., 1975; Werner et al., 1974). Moreover, hyperparathyroidism (PHPT) patients have significantly higher blood glucose levels (Kautzky-Willer et al., 1992) and increased HOMA (homeostatic model assessment) index (Ayturk et al., 2006; Khaleeli et al., 2007; Procopio et al., 2002) compared to controls, a metabolic status that is improved after parathyroidectomy (Akgun & Ertel, 1978; Cheung et al., 1986; Khaleeli et al., 2007, Kautzky-Willer et al., 1992). These observations suggest that the parathyroid hormonal system may influence the control of energy metabolism beyond its classical role in mineral homeostasis.

Both PTH and PTHrP signal through the same G protein-coupled receptor, parathyroid hormone receptor 1 (PTH1R), yet differ markedly in their physiological origin and mode of action. PTH is secreted by the parathyroid glands and acts as a circulating endocrine hormone, whereas PTHrP is primarily produced locally and functions in an autocrine or paracrine manner (Vilardaga et al., 2011). Importantly, PTH1R and PTHrP are expressed in the brain and more abundantly in the hypothalamus (Philbrick et al., 1996; Weaver et al., 1995). Moreover, while the parathyroid glands are the only known source of circulating PTH, recent work indicates that circulating PTH can reach the central nervous system and modulate neuronal population (Zhang et al., 2023). Together, these observations raise the possibility that endocrine and paracrine activation of a common receptor may represent an unexplored mechanism by which hypothalamic neurons diversify metabolic responses. Consistent with this idea, it has been reported that malignancy-induced humoral hypercalcemia, a condition characterized by increased levels of PTHrP, causes an increase in the hypothalamic levels of AgRP (Hashimoto et al., 2007; Iguchi et al., 2001), promoting hunger (Gropp et al., 2005; Jain et al., 2000).

Here, we identify PTH1R signaling in AgRP neurons as a neuroendocrine mechanism that independently controls feeding behavior and adaptive metabolic responses. We demonstrate that PTH and PTHrP engage distinct intracellular signaling pathways downstream of the same receptor, thereby selectively regulating feeding, fat oxidation, nutrient partitioning, and neuronal lipophagy. Together, our findings establish the parathyroid hormonal system as an unexpected regulator of hypothalamic metabolic flexibility and provide a conceptual framework for understanding how specialized neurons convert endocrine and paracrine information into distinct physiological outputs. These discoveries may also pave the way for the development of promising therapeutics for metabolic disorders, such as obesity and metabolic syndrome.

## Results

### Arcuate nucleus-borne PTH1R signaling controls energy metabolism

We investigated *Pth1r* and *Pthrp* expression patterns in different brain structures and peripheral organs, including bone (as positive control) which is reported to highly express both transcripts (Philbrick et al., 1996; Weaver et al., 1995). We found that among the brain regions analyzed, both *Pth1r* and *Pthrp* are predominantly expressed in the hypothalamus (HpT) (**Figure 1A**), a major integrative centre of metabolic feedback regulating energy balance. Within the HpT, we identified by immunofluorescence a high presence of both proteins in the arcuate nucleus (ARC) (**Figure 1B**), localized in the ventral part of the medio-basal HpT adjacent to the third ventricle (3V). Interestingly, PTH1R is distributed all along the rostro-caudal axis, when PTHrP is mainly expressed in the rostral part of the ARC nuclei (**Figure S1A**). Of note, we also detected the presence of PTH1R in cells of the median eminence and along the walls of the 3V (**Figure 1B**). By contrast and as expected, *Pth* is not expressed in any brain areas but exclusively produced by the periphery (i.e parathyroid glands) (**Figure 1A**).

**Figure 1:**
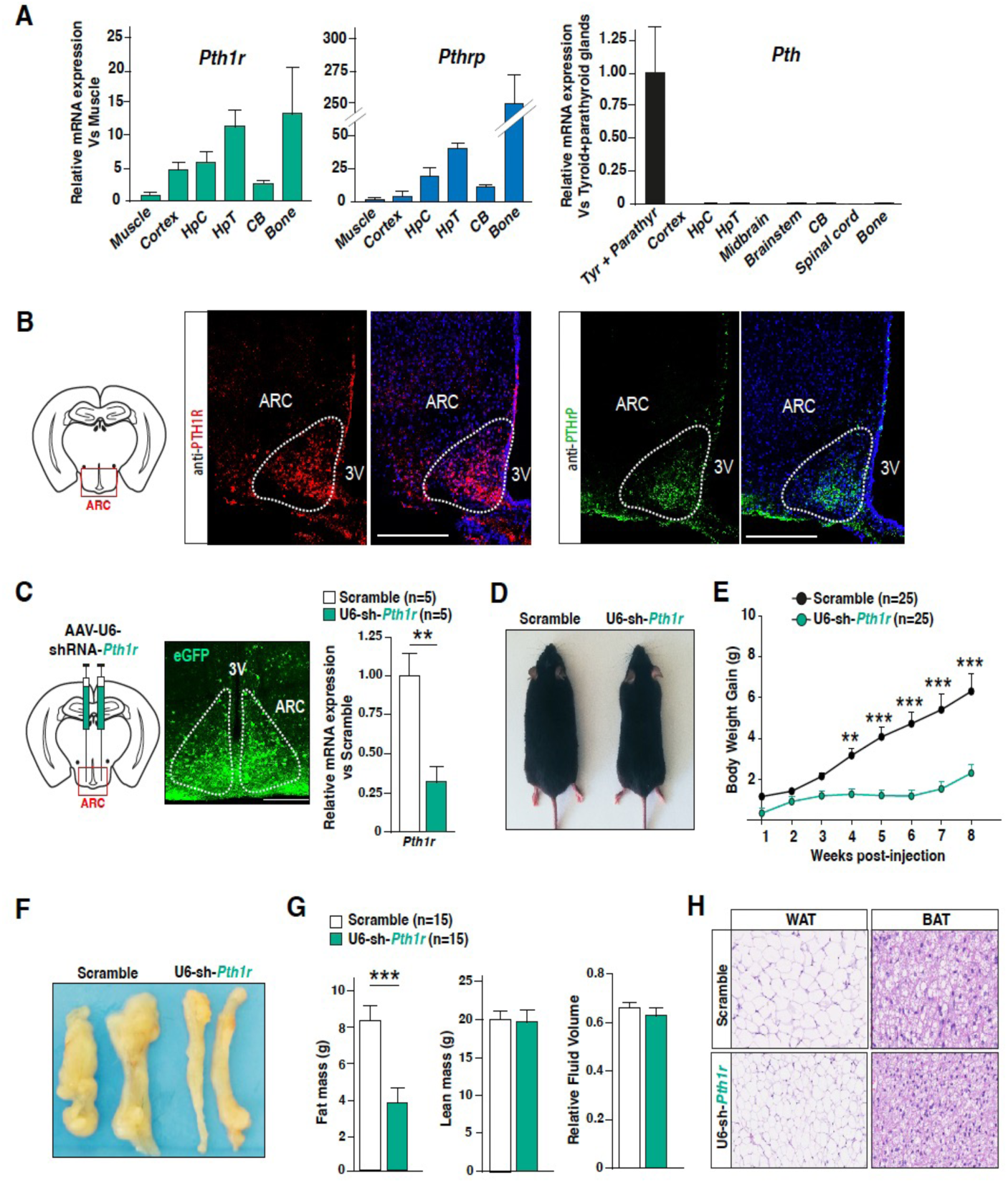
ARC-borne PTH1R contributes to the central control of energy metabolism. **A**) *Pth1r* and *Pthrp* relative gene expression (RT-qPCR performed in triplicate for each sample) in muscle, bone and various parts of the brain [cortex, hippocampus (HpC), hypothalamus (HpT), and cerebellum (CB)] of 3-month-old (3Mo) WT mice (n = 5). *Pth* relative gene expression (RT-qPCR performed in triplicate for each sample) in muscle, liver, kidney, bone, fat, lung, thyroid + parathyroid glands and various parts of the brain [cortex, hippocampus (HpC), hypothalamus (HpT), and cerebellum (CB)] of 3-month-old (3Mo) WT mice (n=3). Quantification of mRNA expression is relative to muscle. **B**) Representative pictures of immunofluorescence staining performed on brain cross-sections of 3 Mo mice, showing that both PTH1R (red) and PTHrP (green) are abundantly present in the ARC (Bregma-2.9). Scale bars, 100 μm. **C**) Schematic representation of the stereotactic injections of viral vectors in the ARC (X=-1.49; Y= +/-0.40; and Z=-5.67), fluorescent microscopy images of brain cross-sections collected 3 weeks after injections (the AAV expresses eGFP, which allows for verification of the site and efficacy of the infection) and bar graphs representing the relative mRNA expression of mice stereotactically injected with AAV-U6-shRNA-Scramble (n=5) or AAV-U6-shRNA-*Pth1r* (n=5). **D-E**) Representative images (**D**) and body weight gain (**E**) of mice stereotactically injected with AAV-U6-shRNA-*Pth1r* (n=25) or AAV-U6-shRNA-Scramble (n=25) in the ARC. **F**) Representative images of mice injected with AAV-U6-shRNA-Pth1r or AAV-U6-shRNA-Scramble fat pad. **G**) Body composition measurements (by NMR analyzer) assessing body fat mass, lean mass and relative fluid volume of mice injected with either AAV-U6-shRNA-*Pth1r* (n=15) or AAV-U6-shRNA-Scramble (n=15). **H)** Representative images of adipocytes in white (WAT) and brown (BAT) adipose tissues of mice injected with the two different experimental groups. Data are expressed as mean ± s.e.m. For overall analysis of metabolic cage parameters, the Mann–Whitney test was used. For body weight analysis, and dark-light analysis of metabolic parameters, the two-way ANOVA was used followed by a Sidak post-hoc test. For fatty acid oxidation, the two-way ANOVA was used with Geisser-Greenhouse correction. \**P* < 0.05, \*\**P* < 0.01, \*\*\**P* < 0.001.

In order to explore if and how PTH1R signaling in the ARC could impact energy metabolism, we downregulated *Pth1r* expression in the ARC, via bilateral stereotactic injection of an adeno-associated virus (AAV) expressing short-hairpin RNA (shRNA) against *Pth1r* under the control of the ubiquitous U6 promoter (**Figure 1C**). A decrease of up to 70% in hypothalamic *Pth1r* expression was validated in the HpT 3 weeks after injection (**Figure 1C**). At a metabolic level, we found that selective *Pth1r* downregulation in the ARC led to a decrease in body weight gain in mice fed with normal chow (**Figure 1D-E**), associated with a decrease in white fat pad size (**Figure 1F**), whole-body fat mass (as assessed by nuclear magnetic resonance spectroscopy) (**Figure 1G**), and lipid vacuole size in both white and brown adipose tissues (**Figure 1H**). Altogether these findings indicate that PTH1R signaling integrity in the ARC is required for optimal control of energy metabolism.

### PTH1R and PTHrP are present in AgRP neurons

Given the functional heterogeneity of the hypothalamic neurons (Cone, 2005), we next assessed the cellular specificity of PTH1R in the HpT. Immunofluorescence analyses using transgenic mice expressing the reporter td-Tomato in AgRP neurons (*Agrp*^Cre:tdTomato^) or POMC neurons (*Pomc*^Cre:tdTomato^) mice revealed a enrichment of PTH1R and PTHrP within AgRP neurons (**Figure 2A**), whereas only sparse labeling was observed in POMC neurons (**Figure S2A**). This observation was also confirmed using AgRP peptide immunodetection with a validated antibody against AgRP (**Figure S2B-C**), combined to *in situ* hybridization (RNAscope) with either *Pth1r* or *Pthrp* probes (**Figure 2B**). Of note, we also detected the presence of PTH1R in cells of the median eminence and along the walls of the 3V (**Figure 1B** and **Figure S1A, S1B**).

**Figure 2:**
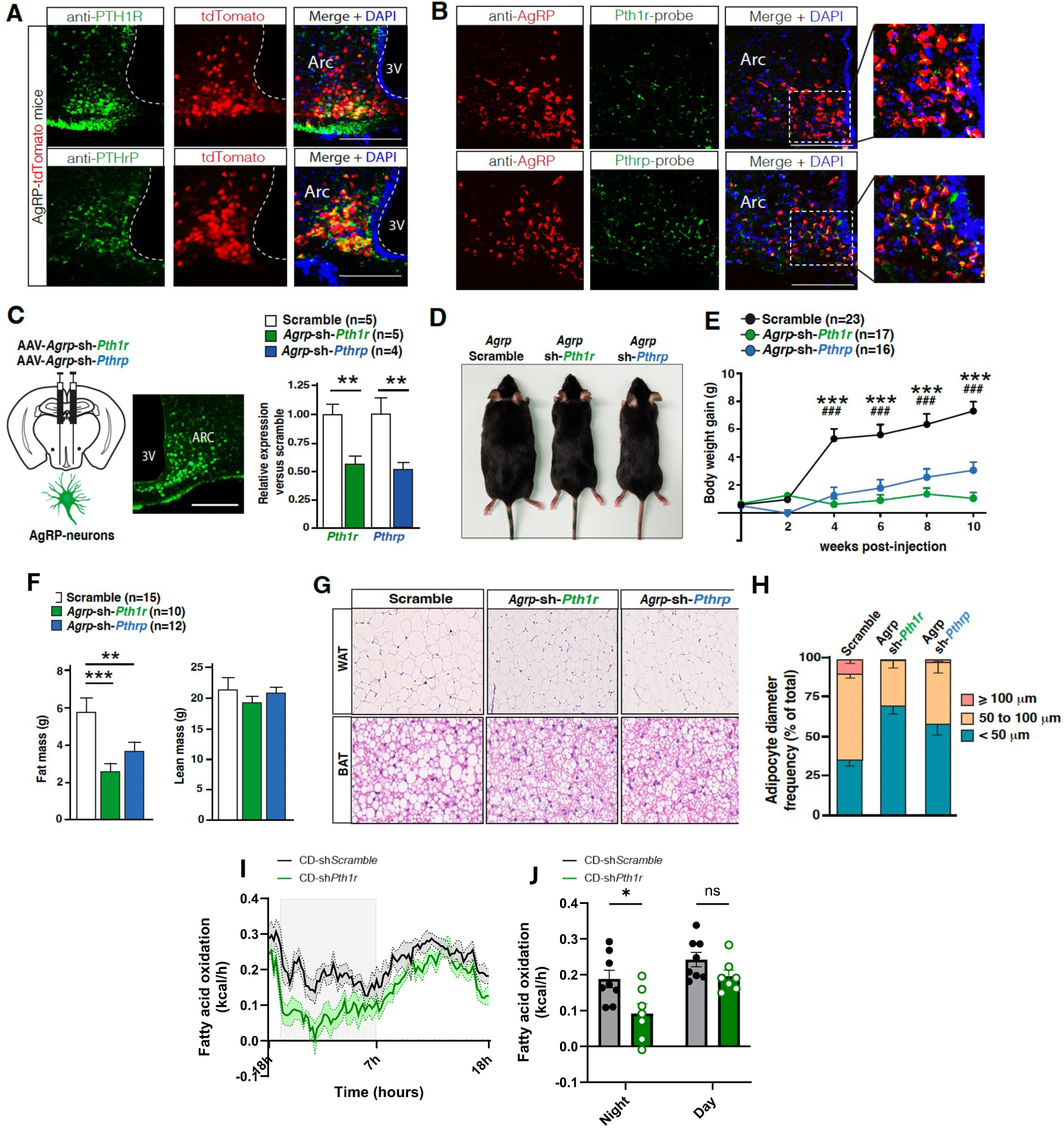
***Pth1r* is expressed in AgRP+ neurons and influences feeding behavior, energy expenditure and fat metabolism. A)** Representative image of immunofluorescence staining depicting the localization of PTH1R and PTHrP (in green) in AgRP neurons (red). The immunostainings were performed on brain cross-sections of 3Mo *Agrp*^Cre:tdTomato^ mice. Scale bars: 100µm. **B)** *In situ* hybridization with immunofluorescent staining showing the expression of *Pthrp* and *Pth1r* (in green) in AgRP (red) neurons. **C**) Schematic representation of the stereotactic injections of viral vectors in the ARC (X=-1.48; Y= +/-0.40; and Z=-5.67), fluorescent microscopy images of brain cross-sections collected 5 weeks after injections (the AAV expresses eGFP under the control of the *Agrp* promoter, which allows for verification of the site of the infection). **D-E**) Representative images (**D**) and body weight gain (**E**) of mice injected with AAV-*Agrp*-shRNA-*Pth1r* (n=17), AAV-*Agrp*-shRNA-*Pthrp* (n=16) and AAV-*Agrp*-shRNA-Scramble (n=23) in the ARC. **F**) Body composition measurements (by NMR analyzer). **G)** Representative images of adipocytes in white (WAT) and brown (BAT) adipose tissues of mice injected with either AAV-*Agrp*-shRNA-*Pth1r* (n=5) AAV*-Agrp*-shRNA-*Pthrp* (n=9) or Scramble (n=14). **H**) Bar plot depicting the frequency of small (<50 µm), big (50–100 µm) and large (≥100 µm) adipocytes in the WAT of mice, 3 weeks after injections of AAVs carrying shRNA against Pthrp or Pth1r (shRNA driven by the *Agrp* promoter). **I-J**) Bar graphs representing (overall mean and day-night mean) whole-body fatty acid oxidation (**J**) of AAV-*Agrp*-shRNA-*Pth1r* (n=7) and Scramble mice (n=8), measured in automated metabolic cages over 5 days, following 2 days of habituation. Data are expressed as mean ± s.e.m. For overall analysis of metabolic cage parameters and WAT adipocytes diameter, one-way ANOVA was used followed by post-hoc Tuckey’s test. For body weight analysis and dark-light analysis of metabolic parameters, the two-way ANOVA was used followed by a Sidak post-hoc test. For fatty acid oxidation, the two-way ANOVA was used with Geisser-Greenhouse correction. \**P* < 0.05, \*\**P* < 0.01, \*\*\**P* < 0.001.

### PTH1R signaling onto AgRP neurons modulates energy balance and fat metabolism

Since modulation of AgRP neuron activity is known to regulate feeding, energy expenditure and peripheral nutrient partitioning (Cavalcanti-de-Albuquerque et al. 2019; Joly Amado et al., 2012; Manceau et al., 2020; Nogueiras et al., 2007; Reichenbach et al., 2018a and b), we next explored more directly the impact of PTH1R signaling on the regulation of AgRP neuron output. Mice were stereotactically injected in the ARC with an AAV expressing shRNA for either Scramble or *Pth1r* under the control of the *Agrp* promoter (**Figure 2C**). The downregulation of *Pth1r* was validated using co-immunofluorescence on HpT brain sections (**Figure S2D**). Then, the mice were subjected to systematic metabolic efficiency phenotyping. We found that mice in which *Pth1r* is specifically downregulated in AgRP neurons (*Agrp*-shRNA-*Pth1r* mice) gained significantly less body weight on standard diet compared to scramble-injected mice (**Figure 2D-E**). Body composition analysis showed a diminution of fat mass in *Agrp*-shRNA-*Pth1r,* without changes in lean mass (**Figure 2F**). Histological examination of BAT and WAT tissues showed that AgRP-specific downregulation of *Pth1r* induced adipocyte shrinkage, caused by a decrease in lipid vesicle size (**Figure 2G-H**). Metabolic efficiency characterization using indirect calorimetry revealed that AgRP-neuron restricted downregulation of *Pth1r* led to a significant decrease in fatty acid oxidation (**Figure 2I-J**) while no differences were observed in neither global energy expenditure (EE), RER, VO2 consumption nor VCO2 production (**Figure S2G**). Importantly, while *Pth1r* knockdown in AgRP neuron invariably resulted in decreased fat oxidation and body weight gain after surgery, cumulative feeding on chow diet was either only marginally affected (see **Figure S2E**) or decreased (see **Figure S2F**) suggesting that change in calorie intake *per se* mihght not the primary mechanism driving body weight change.

Noteworthily, AgRP restricted knockdown of *Pth1r* recapitulated the metabolic consequence of a more general knockdown achieved by stereotactic injection of AAV-U6-*Pth1r*-shRNA observed in **Figure 1C-E**. Altogether, these data suggest that the behavioral and metabolic consequences arising from PTH1R activation in the HpT are relayed by AgRP-neurons.

### Circulating PTH and local PTHrP influence the activity of ARC neurons by modulating PTH1R signaling

The defects in energy balance observed following downregulation of *Pth1r* in AgRP neurons raised the question regarding the specific mechanisms conveying its effects. PTH1R can be potentially activated by two ligands, the circulating PTH, which has been reported to cross the blood-brain barrier (Bühler at al 1997; Joborn et al., 1991; Lourida et al., 2015), and the paracrine/autocrine signal PTHrP, which we showed here to be abundantly present in the ARC nuclei (**Figure 1B, 2A-B** and **S1A**).

Using stereotactic injections of AAV-U6-*Pthrp*-shRNA (general promoter) or AAV-AgRP-*Pthrp*-shRNA (AgRP promoter), we found that the genetic manipulation of its bona fide paracrine ligand *Pthrp* in the ARC cannot fully recapitulate the metabolic consequences observed following downregulation of *Pth1r*. Indeed, we show that *Pthrp* downregulation in the ARC (either in the global ARC area or selectively in AgRP neurons), mimicked the decrease in body weight gain, fat mass in mice downregulated for *Pth1r* selectively in AgRP neurons, but displayed a significant changes in food intake with no changes in energy expenditure (**Figure S3A-D**). We next showed that while systemic injection of PTHrP peptide (100mg/kg BW, as previously described in Kir et al., 2014) did not affect feeding (**Figure 3A**), intracerebroventricular (i.c.v., 40ng/hr) administration of PTHrP (1-34) led to an increase in food intake with no changes in fat oxidation (**Figure 3B**) or other metabolic parameters (**Figure S3E**). Importantly, this effect was abolished by selective *Pth1r* downregulation in AgRP neurons (**Figure 3C**). The ability of PTHrP to act centrally was confirmed by recording of AgRP activity upon PTHrP stimulation. Using fresh brain slices prepared from transgenic *Agrp*^Cre:tdTomato^ mice killed at the beginning of the dark period in a calorie-deficient state, we found that bath-applied PTHrP (100 nM) increased firing rate by 87% in approximately 90% of active *Agrp*^Cre:tdTomato^ neurons (n = 12 out of 13 neurons; basal: 1.72±0.46 Hz versus PTHrP: 3.21±0.75 Hz) (**Figure 3D and Figure S3G**). Most of PTHrP-stimulated *Agrp*^Cre:tdTomato^ neurons returned to their basal firing rate 10 minutes after PTHrP application (i.e. during wash-out) (**Figure S3G**). Thus, central PTHrP can boost orexigenic AgRP neurons during low energy state.

**Figure 3:**
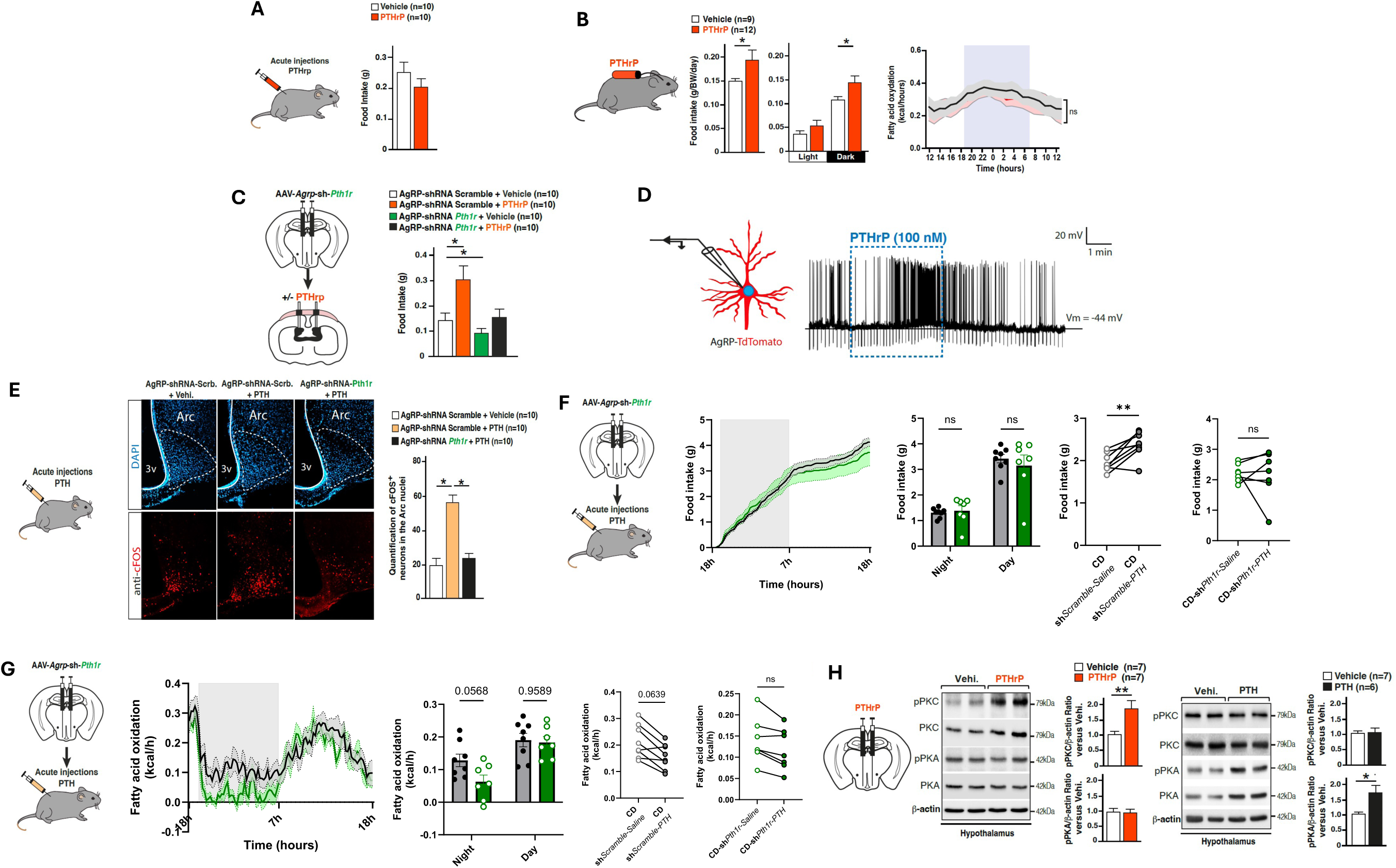
Distinct roles of PTH and PTHrP signaling in the ARC. **A**) Food intake measurement of mice after intraperitoneal injection of PTHrP (1 mg/kg) (n=10) or vehicle (NaCl 0.9%) (n=10). **B**) Food intake measurement of mice injected i.c.v. with PTHrP (40ng) (n=10) or vehicle (NaCl 0,9%) (n=10). **C**) Food intake measurement of mice stereotactically injected with AAV-*Agrp*-shRNA-*Pth1r* (n=10) or Scramble (n=10), and injected i.c.v. 5 weeks later with PTHrP (40ng) (n=10) or vehicle (NaCl 0,9%) (n=10). **D**) Electrophysiological analysis of the acute effect of either PTHrP on AgRP neuronal activity, using patch-clamp recording on brain slices of transgenic *Agrp*^Cre:tdTomato^ mice. Representative cell-attached recording of a *Agrp*^Cre:tdTomato^ neuron was performed before, during and after the application of either PTHrP (100 nM). The duration of the treatment is represented within the blue dashed box. The measurements were performed on n=13 independent neurons from n=8 *Agrp*^Cre:tdTomato^ mice upon PTHrP. **E**) Immunofluorescence staining and quantification of cFOS+ (red) neurons in the ARC, 1 H after intraperitoneal injections with PTH (1 mg/kg) or vehicle (NaCl 0.9%). These injections have been performed in mice priorly stereotactically injected in the HpT with either AAV-*Agrp-*-shRNA-*Pth1r* (n=10) or Scramble (n=10). **F**) Food intake measurement of mice stereotactically injected with AAV-*Agrp*-shRNA-*Pth1r* (n=10) or Scramble (n=10), and injected i.c.v. 5 weeks later with PTH (40ng) (n=10) or vehicle (NaCl 0,9%) (n=10). **G**) whole-body fatty acid oxidation measurement of mice stereotactically injected with AAV-*Agrp*-shRNA-*Pth1r* (n=10) or Scramble (n=10), and injected i.c.v. 5 weeks later with PTHrP (40ng) (n=10) or vehicle (NaCl 0,9%) (n=10). **H**) Representative Western blot and bar graph showing relative quantification of pPKC, PKC, pPKA, PKA proteins in the hypothalamus of mice injected with PTHrP (40 ng/side) (n=7) or vehicle (NaCl 0,9%) (n=7) or injected with PTH (40 ng/side) (n=6) or vehicle (NaCl 0,9%) (n=6). Data are expressed as mean ± s.e.m. For food intake measurement after PTHrP peripheral injection and western blot analysis, the Mann-Whitney test was used. For food intake measurement after PTHrP central injection and cFOS quantification, the one-way ANOVA follow by post-hoc Tuckey’s test was used. For energy expenditure analysis, two-way ANOVA was used followed by a Sidak post-hoc test. \**P* < 0.05, \*\**P* < 0.01, \*\*\**P* < 0.001.

Exploring the role of PTH in the ARC, we showed that intraperitoneal PTH administration is sufficient to induce an increase in cFOS protein expression in the ARC (**Figure 3E**) and that this induction is abolished in AAV-AgRP-*shPth1r*-injected mice, supporting a role of the systemic PTH-PTH1R coupling in AgRP neuron activation. We further demonstrated that intraperitoneal PTH (100mg/kg BW) administration fosters appetite (**Figure 3F**), but also and by contrast to PTHrP, is sufficient to influence fat oxidation neurons (**Figure 3G**). Both effects of PTH in the ARC required, at least in part, the integrity of PTH1R onto AgRP-neurons (**Figure 3G**). Other parameters such RER, EE, water intake, VO2 consumption, VCO2 production and ambulatory activity, remained unchanged after PTH chronic injections (**Figure S3F**). Taken together, these observations indicate the role of circulating PTH in the control of appetite and fat oxidation, while the paracrine/autocrine effects of HpT-borne PTHrP is restricted to the control of feeding. Obviously, the distinct effects of these two ligands rely, at least in part, on PTH1R in AgRP neurons.

### PTH and PTHrP stimulate different PTH1R-associated signaling pathways in the ARC

In the light of our results, one important question that arises is how the same receptor (i.e PTH1R) could regulate different metabolic functions dependently on its activation by either PTH or PTHrP in the HpT. Using biochemical analysis of molecular responses triggered following intra ARC injections of either PTHrP (40ng/hemisphere) or PTH (40ng/hemisphere), we found that PTHrP treatment activates the phospholipase C/inositol trisphosphate/Ca^2+^/protein kinase C (PKC) pathway, while PTH activates the adenylyl cyclase/cAMP/protein kinase A (PKA) pathway (**Figure 3H).** These results suggest that these ligands could modulate AgRP neuron activity by harnessing different GPCR associated signaling pathways. Altogether, these results suggest that binding of either PTHrP or PTH onto PTH1R in the ARC exert discriminatory actions at an intracellular level, engaging distinct signaling pathways and onto the various branches of energy homeostasis that depend on AgRP neuron activity (i.e feeding, energy expenditure or peripheral substrate utilization).

### Hypothalamic PTHrP/PTH1R coupling regulates AgRP levels and fasting-induced re-feeding

We then assessed whether PTHrP signaling is regulated in the hypothalamus following drastic deviation of the energy balance, such as that induced by food deprivation, a situation known to upregulate AgRP levels (Cone, 2005). To address this, we measured the levels of PTH1R and PTHrP in fasted (F), non-fasted (NF) and fasted mice that were re-fed (F-RF) (**Figure 4A**). We observed an increase in AgRP levels that was attenuated following 2 hours of re-feeding (**Figure 4B**), consistently with the known literature upon AgRP regulation (Takahashi & Cone, 2005). Interestingly, the same pattern of protein accumulation was observed with PTHrP and PTH1R, increasing after fasting and returning to normal upon re-feeding (**Figure 4C**). To further validate the link between PTHrP and AgRP expression, we delivered PTHrP in the ARC and found that it promptly increased AgRP levels in the HpT, 1 hour post-injection (**Figure 4D**). Similar results were obtained using either primary HpT neurons or mouse AgRP cell lines (N38) after PTHrP treatment, while no modifications were observed upon PTH treatment (**Figure S4A-C**). Conversely, AgRP-specific downregulation of either *Pth1r* or *Pthrp* in the ARC decreased AgRP protein levels in the HpT (**Figure 4E**). Finally, we assessed whether the integrity of PTH1R signaling onto AgRP neurons is required for the orexigenic action of these neurons in the condition of nutriment scarcity. Feeding response to fasting was assessed in mice with AgRP-restricted knockdown of *Pth1r* or *Pthrp* achieved by ARC-specific delivery of either AAV-*Agrp*-shRNA-*Pthrp* or AAV-*Agrp*-shRNA-*Pth1r.* As shown in **Figure 4F**, feeding response after a fast markedly reduced in *Agrp*-shRNA-*Pthrp* and *Agrp*-shRNA-*Pth1r* mice compared to control mice. Furthermore, we observed a failure in *Agrp*-shRNA-*Pthrp* mice to upregulate hypothalamic AgRP levels in response to fasting (**Figure 4G**) suggesting that PTH1R signaling is an integral part of the adaptive response of AgRP neurons to starvation.

**Figure 4:**
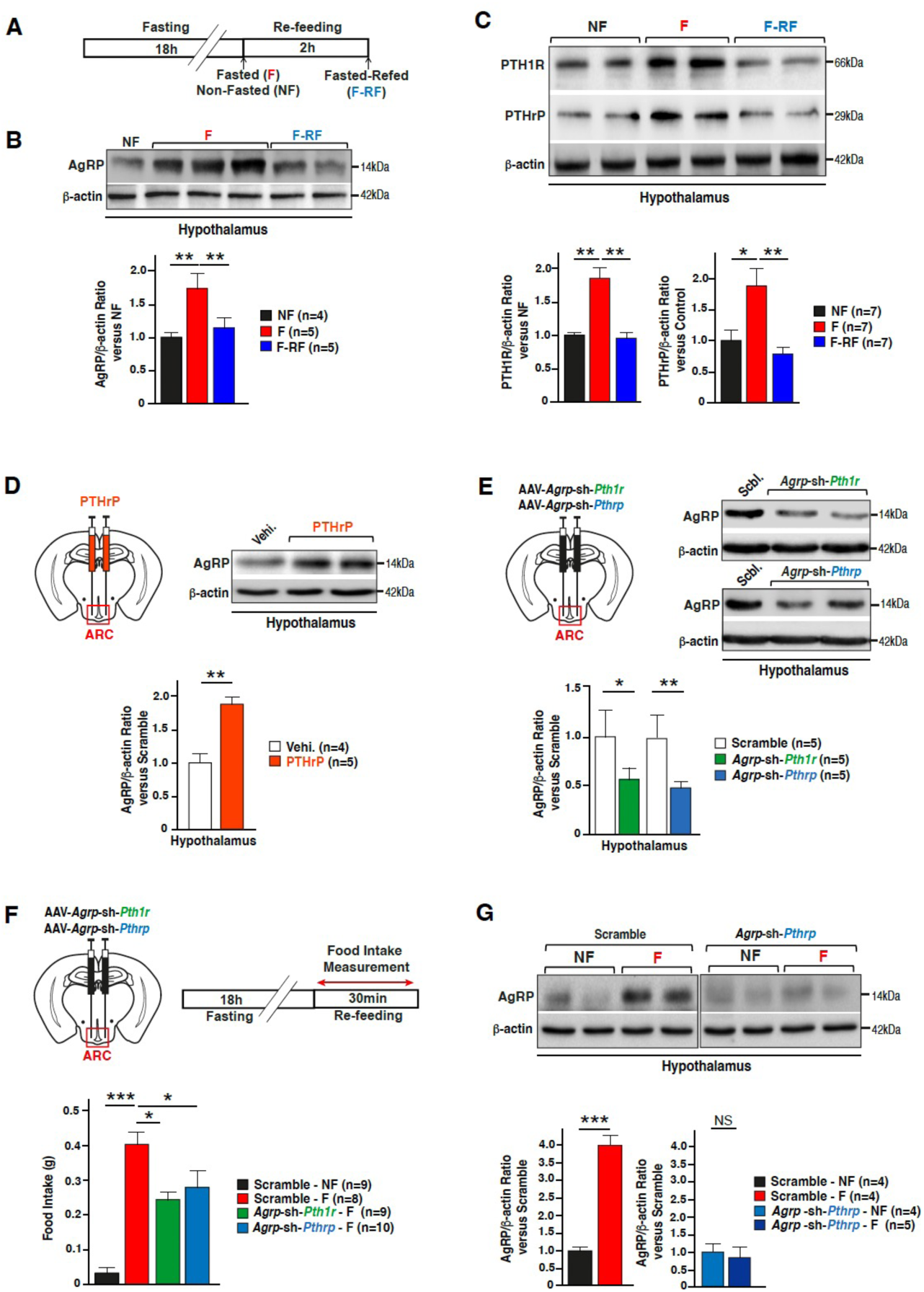
The PTHrP/PTH1r system influences fasting responses by regulating AgRP neuropeptide levels. **A**) Timetable of the fasting paradigm used to assess protein levels of AgRP, PTH1R and PTHrP in the HpT. **B-C**) Representative western blot and bar graph showing relative quantification of AgRP (**B**), PTHrP and PTH1R (**C**) in the hypothalamus of non-fasted (NF), fasted (F) and fasted and refed (F-RF) mice. **D-E**) Representative western blot and bar graph showing relative quantification of AgRP in the hypothalamus of mice stereotactically injected with PTHrP (40 ng / side) (n=5) or vehicle (NaCl 0,9%) (n=4) (**D**) or injected with AAV-*Agrp*-shRNA-*Pthrp* (n=5), AAV-*Agrp*-shRNA-*Pth1r* (n=5) or AAV-*Agrp*-shRNA-Scramble (n=5) (**E**). **F**) Bar graph representing 30 min food intake of NF mice previously injected with AAV-*Agrp*-shRNA-Scramble (n=9) or 18 H-fasted mice previously stereotactically injected with AAV-*Agrp*-shRNA-Scramble (n=8), AAV-*Agrp*-shRNA-*Pth1r* (n=9), AAV-*Agrp*-shRNA-*Pthrp* (n=10). **G**) Representative western blot and bar graph showing relative quantification of AgRP in hypothalamus of mice stereotactically injected with AAV-*Agrp*-shRNA-Scramble, F for 18 H (n=4) or NF (n=4) or injected with AAV-*Agrp*-shRNA-*Pthrp*, F for 16H (n=5) or NF (n=4). Data are expressed as mean ± s.e.m. For western blot quantification, Mann-Whitney test or one-way ANOVA followed by Tuckey’s post-hoc test was used. For food intake measurement, one-way ANOVA followed by Tuckey’s post-hoc test was used. \**P* < 0.05, \*\**P* < 0.01, \*\*\**P* < 0.001.

### PTHrP induces starvation-like hypothalamic lipid degradation

In light of these results, we sought to determine the cellular mechanisms that would underlie these effects. We analysed by liquid chromatography-tandem mass spectrometry (LC-MS/MS) the metabolomic profile of mouse HpT (**Figure 5A-B**) or primary HpT neurons (**Figure S5A-D**) treated with PTHrP. We observed an increase in free fatty acids (FFAs) upon PTHrP compared to vehicle treatments (**Figure 5A and Figure S5A**), reflecting an induction of lipid droplet degradation and increase in FFA availability. Our PCA biplot also indicated that vehicle and PTHrP datasets were orthogonal to each other, indicating an increase in the proportion of FFAs by PTHrP (**Figure 5B and Figure S5B-D**).

**Figure 5:**
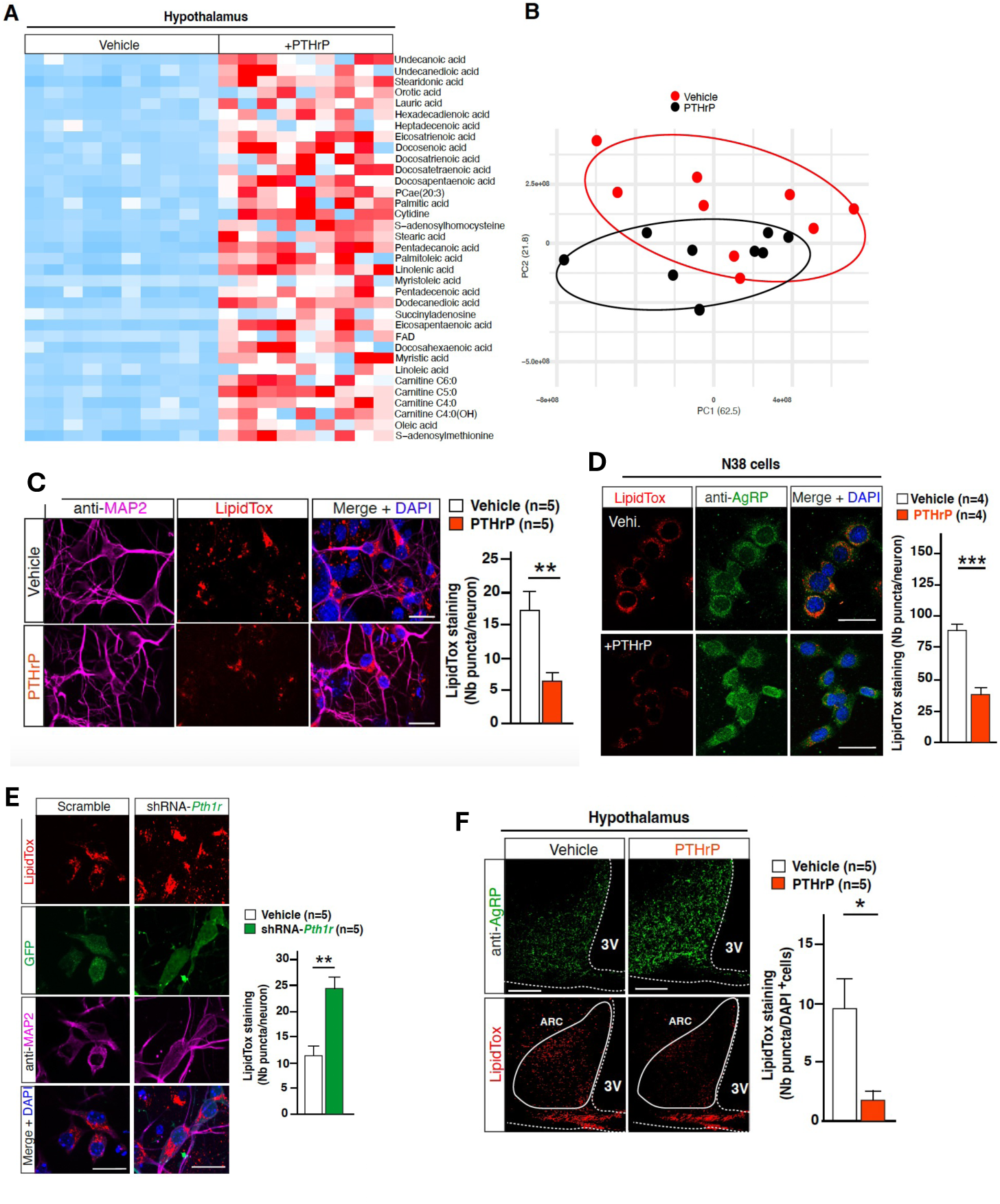
PTHrP promotes the degradation of intrinsic lipids and the production of free fatty acids in the HpT. **A**) Heatmap representing free fatty acid relative abundance in the hypothalamus of vehicle-and PTHrP-treated mice. Analysis was performed using liquid chromatography-tandem mass spectrometry (LC-MS/MS). Representation of the metabolites was generated using the heatmap function in R. **B**) PCA biplot showing that the datasets collected after treatment with either vehicle or PTHrP are orthogonal to each other. **C**) Representative images and quantification of LipidTox staining (red) in MAP2 positive cells (purple) and GFP positive cells (green) in primary hypothalamic neurons infected with Lenti-shRNA-*Pth1r* (n=5) or Lenti-shRNA-Scramble (n=5) expressing GFP. Quantification was made on ≥ 15 replicates. **D**) Representative images and quantification of LipidTox staining (red) in AgRP positive neurons (green) in hypothalamic N38 cells treated with PTHrP (100 nM) (n=5) or Vehicle (n=5). **E**) Representative images and quantification of LipidTox (red) puncta and AgRP (green) fluorescence intensity in N38 cells treated with TCS21311 (PKC inhibitor, 75 µM). Quantification was made on ≥ 15 replicates. **F**) Representative images and quantification of LipidTox (red) puncta in AgRP (green) neurons in hypothalamus of mice stereotactically injected with PTHrP (40 ng/side) (n=5) or vehicle (NaCl 0,9%) (n=5). Quantification was made on ≥ 15 replicates. Data are expressed as mean ± s.e.m. For puncta counting and fluorescence intensity analysis, Mann-Whitney test or one-way ANOVA followed by Tuckey’s post-hoc test was used. \**P* < 0.05, \*\**P* < 0.01, \*\*\**P* < 0.001.

Fasting increases the levels of circulating FFAs that are rapidly taken up by organs and stored in the form of lipid droplets, mainly composed by triglycerides, until lipolytic mechanisms then break them down to provide energy. Importantly, intra-neuronal FFA availability has been previously described as important in the regulation of fasting responses (Kaushik et al., 2011). In view of our observations, we used LipidTox, a neutral lipid dye, to measure the amount of lipid droplets in either primary HpT neurons (**Figure 5C**) or N38 cell line (AgRP-like neurons) (**Figure 5D**) treated with PTHrP. In both experimental sets, we observed a massive reduction in the amount of lipid droplets, while lipid droplet accumulation was observed in primary cells with selective *Pth1r* downregulation (**Figure 5E**). Consistent with the notion of a segregated action of PTH and PTHrP onto AgRP neurons, PTH treatment did not modulate lipid droplets degradation, suggesting alternative mechanisms for the action of this ligand (**Figure S5E**). Of note, this effect on lipid droplets degradation is associated to an increase in AgRP levels in the ARC nuclei (**Figure 5D**). Lastly, the effect of PTHrP on lipid droplets degradation, was also assessed *in vivo,* where acute PTHrP administration in the ARC promptly increased AgRP levels and decreased the amount of LipidTox staining in the ARC (**Figure 5F**). Collectively, these findings suggest that the paracrine PTHrP/PTH1R system displays unique feature as compared to PTH activation, and selectively modulates HpT FFA availability in AgRP neurons.

### PTHrP-induced lipid degradation is driven by the autophagy machinery

Lipid degradation can be the result of either enzyme-driven lipolysis in the cytosol or lipophagy. Accumulating evidence highlights the role of this selective form of autophagy in nutrient-deficient situations in various tissues, including in the HpT (Kaushik et al., 2011).

To examine whether PTHrP-induced increase in hypothalamic FFA levels could be due to activation of autophagy, we first confirm that PTHrP delivery in the ARC significantly upregulates key autophagy proteins, such as ATG5/12 and LC3II, together with AgRP in the HpT (**Figure 6A**). By contrast, PTH treatment did not affect the levels of these autophagy proteins markers (**Figure 6B**). To assess more specifically autophagosome (AP) formation and autophagic flux, we used the pH-sensitive tandem mRFP-GFP-LC3 (fusion of mRFP, GFP and LC3) reporter that highlights AP as yellow puncta and autophagolysosomes (post-lysosomal fusion) as red puncta (the acidic environment inside the lysosome quenches the fluorescent signal of GFP). We found that PTHrP induced an increase in the number of both APs and autophagolysosomes (**Figure S6A**), indicative of an increase in both autophagy formation (GFP+/RFP+) and flux (GFP-/RFP+). We then confirmed that PTHrP stimulates the autophagy machinery via its receptor PTH1R, by using PTHrP-treated primary HpT neurons transfected with a lenti-shRNA for *Pth1r*. We observed that while PTHrP treatment decreased p62 levels, *Pth1r* downregulation blocked this effect (**Figure S6B**), meaning that PTH1R is necessary for PTHrP’s effects on autophagic flux in AgRP neurons. Finally, we treated primary HpT neurons with PTHrP and an inhibitor of autophagic flux, Bafilomycin-A1 (BAF), which blocks the autophagosome-lysosome fusion and subsequent protein degradation in lysosomes. We found that BAF abolishes the PTHrP-triggered reduction in intrinsic lipid droplets (**Figure 6C**), suggesting an implication of the autophagy process in PTHrP’s effect on intracellular lipid mobilization. Taken together these results suggest that PTHrP/PTH1R coupling harnesses the autophagy machinery in AgRP neurons towards lipid degradation.

**Figure 6:**
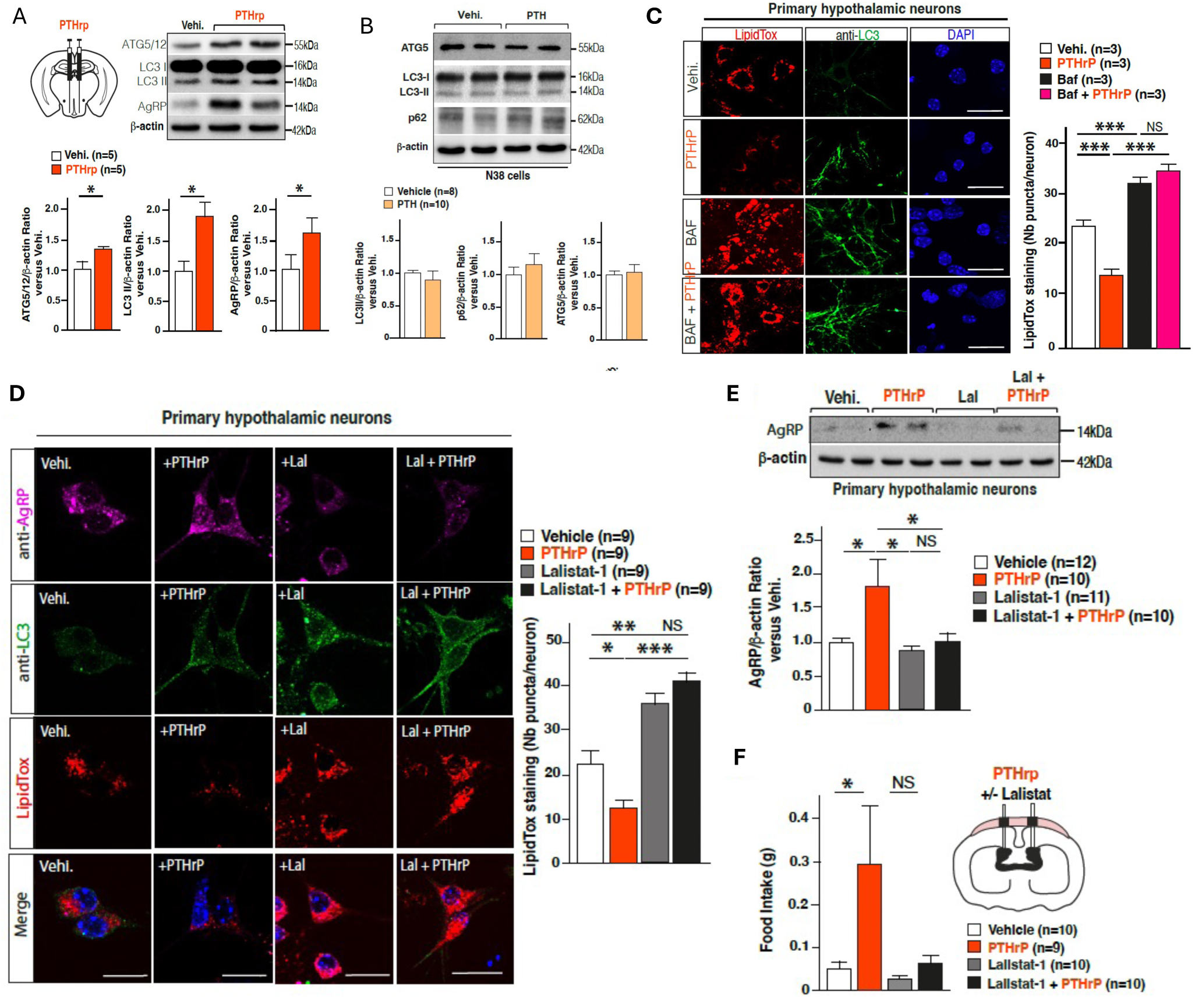
**PTHrP upregulates AgRP levels via a modulation of lipophagy. A-B**) Representative Western blot and bar graph showing relative quantification of the autophagy markers LC3-I and LC3-II, p62 and ATG5-12 in the HpT of mice stereotactically injected with PTHrP (40ng/hemisphere) (n=5), PTH (40ng/hemisphere) (n=5), or vehicle (NaCl 0,9%) (n=5) in the ARC. **C**) LipidTox staining (red) combined with LC3 immunofluorescence staining (green) performed in primary HpT neurons after treatment with either PTHrP (100nM), Bafilomycin (Baf, 1mM) or both for 4 H. Primary HpT neurons were treated at day in vitro (DIV) 12. Quantification was made on n=3 neuronal preparations and the quantification was performed on n ≥ 15 individual HpT neurons per group (each neuronal preparation was derived from at least 12 different embryonic HpT). Scale bar: 20 μm. **D**) Representative images of LipidTox staining combined with immunofluorescence staining for LC3 and AgRP (scale bar: 20 μm) performed on primary HpT neurons treated with either PTHrP (100nM), Lalistat-1 (10mM) or both (puncta quantification was performed on n=3 independent primary cultures). Analysis was performed on n=9 experimental replicates; n ≥ 20 individual neurons per group were used in each replicate. Scale bar: 20 μm. **E**) Representative Western blot and bar graph showing relative quantification of AgRP in primary hypothalamic cells treated with vehicle (n=12), PTHrP (100 nM) (n=10), Lalistat-1 (10 µM) (n=11) or PTHrP + Lalistat-1 (n=10). **F**) Bar graph showing 4 H food intake in mice who received i.c.v. injection of PTHrP (50 ng) (n=10), Lalistat-1 (100 µg) (n=11), PTHrP + Lalistat-1 (n=10) or vehicle (n=12). Lalistat-1 was injected 16 H before the injection of PTHrP and the start of food intake measurement. Results are given as mean ± s.e.m. One-way ANOVA followed by Dunn’s post-hoc and Mann–Whitney tests were used for statistical analysis. \**P* < 0.05, \*\**P* < 0.01, \*\*\**P* < 0.001. NS: non-significant.

### PTHrP-induced lipophagy controls AgRP levels and fasting-induced adaptive responses

To analyze the participation of lipophagy in PTHrP’s effects on intraneuronal lipids, we examined whether inhibition of lipophagy would block the PTHrP-induced lipid droplet degradation in HpT neurons. To that end, we treated HpT neurons with Lalistat-1, a potent and specific competitive inhibitor of lysosomal acid lipase, known to block lipid droplet degradation by the lysosome. Co-treatment of PTHrP with Lalistat-1 hindered PTHrP’s effect on lipid droplet degradation (**Figure 6D**), confirming the implication of lipophagy in PTHrP’s lipolytic effects. To establish a mechanistic link between PTHrP-induced lipophagy and AgRP neuropeptide levels, we measured the amount of AgRP protein in lysates of PTHrP-treated primary HpT neurons after lipophagy inhibition with Lalistat-1. We found that Lalistat-1 blocked the PTHrP-induced increase in AgRP levels (**Figure 6D-E**). At a physiological level, we confirmed that PTHrP-induced feeding is, at least in part, dependent on the lipophagic processes and can be abolished by central blockage of the lipophagic process (**Figure 6F**). These results indicate that PTHrP/PTH1R coupling upregulates AgRP levels, and consequently promotes feeding behavior, by stimulating the lipophagy process in AgRP neurons.

### PTH1R signaling controls the adaptive response of AgRP neurons to circulating orexigenic signals

The gut hormonal factor ghrelin is the only known orexigenic circulating hormone that acts directly on AgRP neurons, inducing rapid transcription of orexigenic neuropeptides, increasing food intake and promoting fat deposition (Nakazato et al., 2001). Ghrelin-induced increase in chow intake has been shown to be dependent of the integrity of AgRP neurons (Chen et al., 2004; Denis et al., 2015; Luquet et al., 2007). We show herein that alteration of PTH1R signaling impaired AgRP neuron-mediated response to a fast, suggesting a convergent mechanism by which ghrelin and PTH1R activation together control AgRP neuron activity. Indeed, systemic injection of ghrelin led to an increase in both PTH1R and PTHrP levels (**Figure 7A-B**), together with a stimulation of the autophagy machinery both *in vitro* and *in vivo* (**Figure 7A-B**). Moreover, similarly to PTHrP action we found that ghrelin can induce lipid droplet degradation in AgRP-like neurons (N38 cell line) and that this induction is prevented by Lalistat-1 treatment (**Figure 7C**). Importantly, selective knockdown of *Pth1r* in AgRP neurons impaired the ghrelin-induced increase in feeding, further confirming the convergence of ghrelin and PTH1R signaling action onto AgRP neurons (**Figure 7D**). These observations suggest that ghrelin could be an upstream regulator of the PTH1R system in AgRP neurons and that PTH1R/PTHrP coupling may act as a modulator of the effects of key systemic metabolic cues in AgRP neurons for the regulation of energy metabolism.

**Figure 7:**
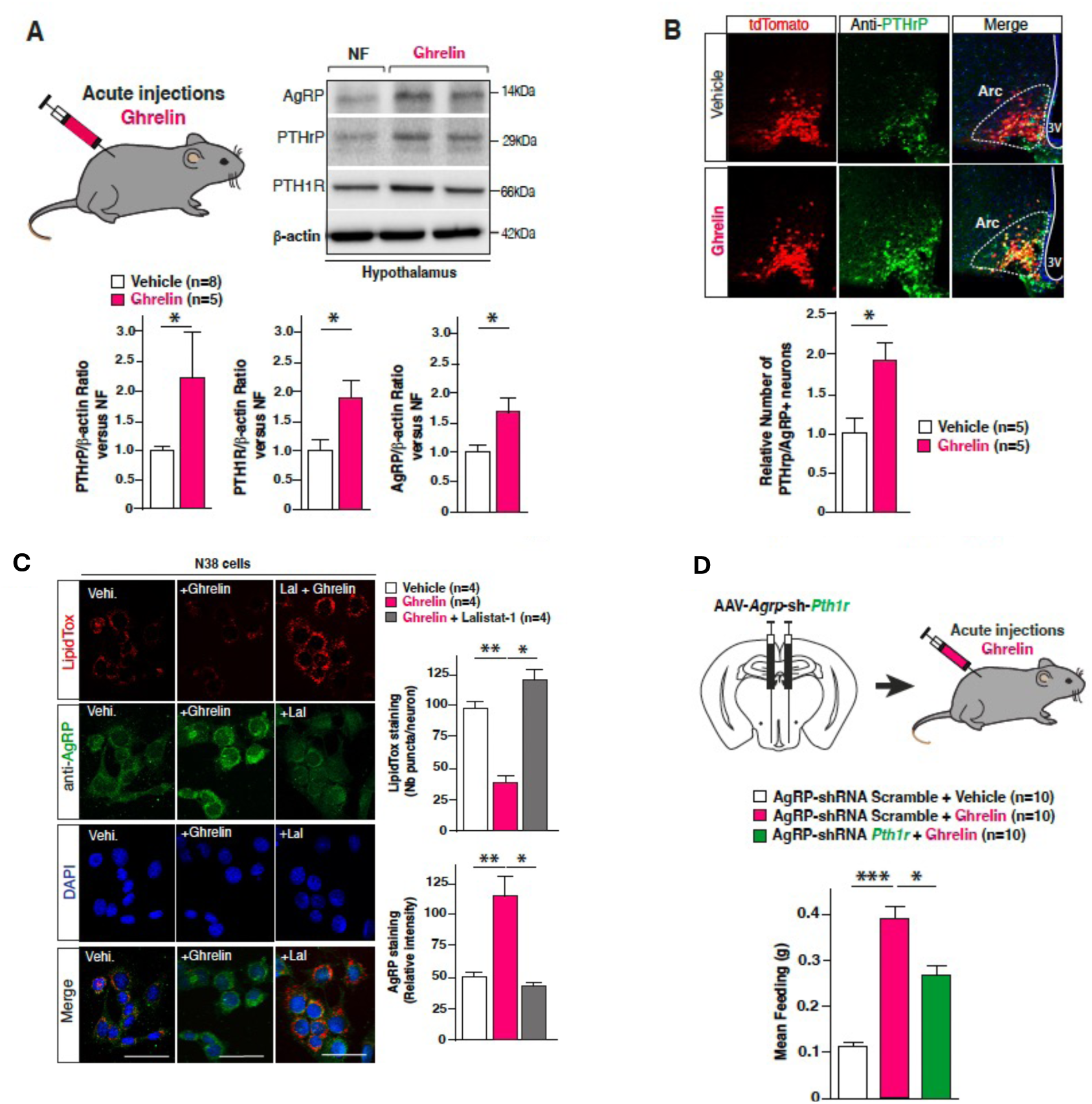
PTHrP is involved in the transmission of feeding behavior induced by ghrelin. **A**) Representative western blot and quantification of AgRP, PTHrP and PTH1R after ghrelin (3 mg/kg, n=5) or vehicle (NaCL 0,9%, n=8) i.p. injection. **B**) Representative pictures of immunofluorescence staining of PTHrP and relative quantification of AgRP positive cells for PTHrP in *Agrp*^Cre:tdTomato^ mice injected with ghrelin (3 mg/kg, n=5) or vehicle (NaCl 0,9%, n=5). **D**) Representative pictures of immunofluorescence staining and quantification of AgRP and LipidTox in N38 cells after treatment with either ghrelin (n=4), ghrelin and lalistat-1 (n=4) or vehicle (NaCl 0,9%) (n=4). **E**) Bar graphs representing food intake in mice stereotactically injected with AAV-*Agrp*-shRNA-*Pth1r* then with ghrelin (3 mg/kg, n=10) 5 weeks later, or injected with AAV-*Agrp*-shRNA-Scramble then ghrelin (n=10) or vehicle 5 weeks later. Data are expressed as mean ± s.e.m. For western blot and PTHrP counting analysis, Mann-Whitney test was used. For LipidTox puncta, AgRP relative intensity and food intake measurement analysis, one-way ANOVA followed by Tuckey’s post-hoc test was used. *P < 0.05, **P < 0.01, ***P < 0.001.

## Discussion

This study identifies the PTH hormonal system as a previously unrecognized signaling module governing hypothalamic control of energy metabolism. We demonstrate that circulating PTH and locally produced PTHrP converge on the same receptor, PTH1R, yet elicit distinct physiological outputs by engaging different intracellular signaling programs within AgRP neurons. By coupling endocrine and paracrine information to specialized cellular mechanisms, the PTH/PTHrP–PTH1R axis independently regulates nutrient partitioning, fatty acid utilization, and adaptive responses to fasting. More broadly, our findings reveal how a single GPCR can diversify the physiological functions of hypothalamic neurons through ligand-specific signaling.

A central feature of energy homeostasis is the ability of hypothalamic neurons to integrate hormonal, nutrient, and local signals. Foundational work on leptin and melanocortin circuits established how POMC and AgRP neurons respond to circulating factors such as ghrelin and insulin, as well as sensory and gut-derived inputs (Manceau et al., 2020). Our findings extend this framework by introducing PTH/PTHrP as an additional regulatory layer, combining endocrine and paracrine modes of action within AgRP neurons.

In line with the emerging connection between bone physiology and energy metabolism (Ducy et al., 2000; Karsenty, 2006), we identify PTH1R as a functional component of hypothalamic circuits. Clinical observations support this link, as circulating PTH and PTHrP levels correlate with obesity, BMI, glucose levels, and insulin resistance (Adam et al., 2010; Bolland et al., 2006; Corbetta et al., 2018; Grey & Reid, 1995; Guasch et al., 2011; Hjelmesæth et al., 2009; Kovesdy & Kalantar-Zadeh, 2007; Martínez-García et al., 2012; Ministrini et al., 2020; Reis et al., 2007; Roca-Rodriguez et al., 2015; Saab et al., 2010; Tran et al., 2014). Moreover, hyperparathyroidism is associated with altered glucose metabolism and insulin resistance, which can be improved following parathyroidectomy (Akgun & Ertel, 1978; Kautzky-Willer et al., 1992; Ayturk et al., 2006; Khaleeli et al., 2007; Procopio et al., 2002; Bannon et al., 1988; Dent, 1962; Ljunghall et al., 1983; Taylor & Khaleeli, 1997; Taylor & Khaleeli, 2001; Walsh et al., 1975; Werner et al., 1974; Cheung et al., 1986). In addition, elevated PTHrP levels in malignancy are associated with increased orexigenic signaling in the ARC (Hashimoto et al., 2007; Iguchi et al., 2001), collectively supporting a role for this system in central metabolic regulation.

Although acting on the same receptor, PTH and PTHrP exert complementary and non-redundant effects. Circulating PTH behaves as a metabolic signal sensed by AgRP neurons to regulate energy expenditure and lipid oxidation without directly affecting feeding behavior. In contrast, PTHrP acts locally to enhance AgRP neuron activity during energy deficit, promoting feeding and adaptive refeeding responses. This distinction is further supported by our observation that AgRP-specific Pthrp downregulation reduces adiposity without altering energy expenditure, highlighting a role in nutrient partitioning and metabolic adaptation.

Mechanistically, these effects rely on ligand-specific GPCR signaling. PTHrP activates the PLC/IP3/Ca²⁺/PKC pathway, driving lipophagy-dependent lipid mobilization and increasing intracellular FFA availability, which supports AgRP expression and feeding. Conversely, PTH preferentially activates the cAMP/PKA pathway. This divergence is consistent with previous reports showing differences in PTH1R signaling dynamics, including sustained intracellular signaling for PTH and membrane-restricted signaling for PTHrP (Cheloha et al., 2015; Nemec et al., 2022).

These findings align with the emerging view that AgRP neurons exert segregated control over feeding, energy expenditure, and substrate utilization through distinct neuronal subpopulations and signaling pathways (Bouyakdan et al., 2022; Betley et al., 2013; Manceau et al., 2020). The differential distribution of PTHrP and PTH1R along the rostro-caudal axis of the ARC may provide a neuroanatomical basis for this functional specialization.

Given the well-established context-dependent effects of PTH in bone, where intermittent and sustained exposures elicit anabolic versus catabolic outcomes, respectively (Silva & Bilezikian, 2015; Datta et al., 2017), it is plausible that distinct temporal dynamics of PTH signaling in the hypothalamus may similarly engage divergent physiological responses. In this context, our experimental paradigm does not address acute PTH action, and we therefore cannot exclude the possibility that rapid or transient PTH signaling may exert additional effects, including the modulation of feeding behavior. Addressing the temporal dimension of PTH signaling in the brain will be important for fully understanding its contribution to energy homeostasis.

Beyond fasting, the PTH/PTHrP–PTH1R system is likely sensitive to broader physiological and environmental challenges. Conditions such as physical exercise, ambient temperature, or other forms of metabolic stress are known to influence energy expenditure and substrate utilization, and may modulate this signaling axis. Investigating these contexts will be essential to fully define the physiological relevance of this system. Such factors may also contribute to variability observed in human clinical studies linking PTH or PTHrP levels to metabolic phenotypes, where environmental and stress-related parameters are often not controlled.

At the cellular level, our data identify PTHrP-induced lipophagy as a key mechanism linking intracellular lipid metabolism to neuronal output. By regulating FFA availability, this pathway supports AgRP neuron activation during nutrient scarcity (Kaushik et al., 2011; Singh et al., 2009; Joly-Amado et al., 2012; Joly-Amado et al., 2014). The convergence between ghrelin and PTH1R signaling further suggests that this system amplifies canonical orexigenic pathways.

In summary, our findings establish PTH1R signaling as a central regulator of energy balance, integrating both circulating (PTH) and local (PTHrP) inputs to generate distinct metabolic outputs in AgRP neurons. This work highlights how ligand-specific GPCR signaling expands the functional repertoire of hypothalamic circuits and provides new insight into the coordination of feeding, energy expenditure, and metabolic flexibility.

### Limitation of the study

In this study we do not provide any insight in the cellular processes that siege at the entry and action of circulating PTH, nor how is orchestrated the potential competition of PTH and PTHrp onto PTH1R. While it is possible that median eminence tanycytes may play a key role in regulating this process, further studies are warranted to fully decipher this mechanism. In addition, our study falls short of providing a full description of the specific projections that could be assigned to subpopulations of AgRP neurons in which PTH or PTHrP action onto PTH1R might engage segregated neural networks.

## Acknowledgements

We thank G. Gangarossa for critical suggestions during the course of the study. We thank the viral production (VVTG), Imaging, Metabolomics, Histology, Behavioral and Metabolism and animal house facility (LEAT) platforms of the SFR Necker. This work was supported by FRM labellisation (F.O., E.S., D.R.G., M.R.), FRM Amorçage (F.O.), MSD-Avenir (F.O., S.L.), Fondation des Gueules Cassées (E.S.), EFSD (E.S. and F.O.), NRJ foundation for Neuroscience (F.O), HFSP (F.O.), ATIP-AVENIR Program (F.O.), Emergence Ville de Paris (F.O.), AGEMED-INSERM (F.O.), INSERM-Transfert (F.O.), MetaboBrain (F.O, S.L) and FSER (F.O).

## STAR Methods

### Animals

Adult male C57BL/6j mice were purchased from Janvier labs (St Berthevin, France). All mice were 3 months of age at the start of experiments. For all experiments, we used littermates as controls. Upon arrival, mice were housed in groups of five in polycarbonate cages (35.5 x 18 x 12.5) and maintained in standard conditions (controlled room temperature and humidity, 12h/12h light/dark cycle, with lights on at 8:00 AM, *ad libitum* access to dry food pellets and water) at the Institut Necker’s animal core facility, officially registered for experimental studies on rodents (Approval number for animal care facilities: A 75-15-01-6 2014-720).

*Pth*^-/-^ (**Pth^tm1a(EUCOMM)Wtsi^**) *and Pthrp^+/-^ (***Pthlh^tm1a(KOMP)Wtsi^**) mice were obtained from EUCOMM program - The Wellcome Trust Sanger Institute Hinxton, Cambridge, UK. The mice were maintained in standard conditions (controlled room temperature and humidity, 12h/12h light/dark cycle, with lights on at 8:00 AM, *ad libitum* access to dry food pellets and water) at the Institut Necker’s animal core facility, officially registered for experimental studies on rodents (Approval number for animal care facilities: A 75-15-01-6 2014-720). The metabolic phenotyping was performed on 3 month-old male mice and compared to control littermates.

*Pomc*^Cre^; *Pth1r*^flox/flox^ *mice* were obtained by crossing *Pomc*^Cre^ (Stock No. 005965;, EUCOMM-EM:07507) with PTH1R^flox/flox^ (generated from **Pth1r^tm1a(EUCOMM)Hmgu^**-EUCOMM program - The Wellcome Trust Sanger Institute Hinxton, Cambridge, UK). The mice were maintained in standard conditions (controlled room temperature and humidity, 12h/12h light/dark cycle, with lights on at 8:00 AM, *ad libitum* access to dry food pellets and water) at the Institut Necker’s animal core facility, officially registered for experimental studies on rodents (Approval number for animal care facilities: A 75-15-01-6 2014-720). Experiments were performed on 3 months-old male mice using PTH1R^flox/flox^ as control littermates.

All animal studies were performed in compliance with the French legislation (Decree of 1 February 2013, 2013-118) and the European Communities Council Directive of 22 September 2010 (2010/63/EEC). All animal experiments were designed according to the 3R’s rules and approved by the French Ministry of Research (A 75-14-083).

### Transgenic Agrp and Pomc reporter mice

Transgenic *Agrp*^Cre:tdTomato^ and *Pomc*^Cre:tdTomato^ mice were obtained by crossing *Agrp*^Cre^ (Stock No. 012899) or *Pomc*^Cre^ (Stock No. 005965) with *tdTomato*^flox/flox^ (Stock No. 007914) mice, which were all purchased from Jackson Laboratories. Experiments were made on 6/9-week old male that were heterozygous for *AgRP* (cre+/−) or *POMC* (cre+/−), and homozygous for *tdTomato*. Animals were housed at 22.5 ± 1°C. Before experiments, animals had free access to standard diet (A04, SAFE) and water. All protocols including animals were reviewed by our local ethic committee in strict accordance with European Community guidelines and agreed by the French Ministry of Higher Education and Research (accreditation No. 00853.01).

### Agrp-ablated mice

Homozygous AgRP^DTR/DTR^ (C57Bl/6 genetic background) were bred with C57Bl/6 wild-type AgRP^+/+^ mice, such that the offspring would be AgRP^DTR/+^ heterozygotes. Heterozygous AgRP^DTR/+^ mice with the <u>Diphtheria Toxin Receptor (DTR)</u> targeted to the AgRP locus have been previously described by (Denis et al., 2015; Joly-Amado et al., 2012; Luquet et al., 2005). Newly born AgRP^DTR/+^ mice, 2-15 days old, received a subcutaneous (interscapular) injection (12.5µl/g) of either DT (Diphtheria Toxin, Sigma D0564-1mg, 10 µg/Kg) or PBS 1X. In consequence, these mice were divided into 2 groups: AgRP-ablated and AgRP-control mice respectively.

### Surgical Procedures

Prior to any surgical procedure, mice were anesthetized by intraperitoneal injection of ketamine hydrochloride (20mg/ml BW; 1000 Virbac) and xylazine (1000mg/ml BW; Rompum 2%; Bayer) and place in a stereotaxic frame (900SL; Kopf Instruments). Coordinates were identified using the Paxinos mouse brain atlas. Ophthalmic eye ointment was applied to the cornea to prevent desiccation during surgery. The area around the incision was trimmed and an iodine solution was applied (Vétédine).

#### Stereotactic injections

For viral stereotactic injections, mice were injected with 1.0 µL virus in each side of the ARC using the following coordinates relative to lambda: (+/-0.4 mm ML,-1.48 mm AP, 5.69 mm DV). The skin was closed with sutures. The virus injected were purchased from Vector Biolabs: AAV9-GFP-*U6*-m-*Pth1r*-shRNAmir, AAV0-GFP-*U6*-m-*Pthrp*-shRNAmir, AAV9-GFP-*U6*-Scramble-shRNAmir, AAV9-*Agrp*-eGFP-*Pth1r*-shRNAmir, AAV9-*Agrp*-eGFP-*Pthrp*-shRNAmir and AAV9-*Agrp*-eGFP-Scramble-shRNAmir. All behavioral experiments or tissue harvest were performed at least 3 weeks after injections.

For drug stereotactic injections, mice were injected in each side of the ARC with 1.0 µL of either PTHrP (40 ng/µL; 4017147; Bachem) or PTH (40 ng/µL; 4011474; Bachem), using the same coordinates as previously described. The entire HpT was carefully dissected 15 minutes or 1 hour after the injections.

#### Canula implantation

For canula implantations, a hole was made at the following coordinates relative to lambda (-1mm ML,-0.5 mm AP). A guide canula with a dummy canula (62003; 26 gauge; RWD) was inserted through the hole into the lateral ventricle and secured to the skull with dental cement (Super-Bond C& B; Sun Medical). Lastly, the scalp incision was sutured around the canula. All infusions were performed 1 week after canula implantations.

#### Osmotic pumps with i.c.v. distribution implantation

For osmotic pumps implantation, the Alzet micro-osmotic pumps (model 1002; Charles River) were subcutaneously installed in the back of the mice and connected to Alzet Brain Infusion Kit (model III) implanted at the following coordinates (-1mm ML,-0.5mm AP). The pumps were filled 24 hours before implantation with freshly prepared PTHrP (40ng/hr; 4017147; Bachem) or PTH (40ng/hr; 4011474; Bachem) dilutions. All behavioral experiments were performed 2 weeks after implantations.

### Indirect calorimetry measurements

Indirect calorimetry was performed using automated metabolic cages (Labmaster, TSE Systems GmbH, Bad Homburg,Germany) where mice are individually housed for consecutive days and which include bedding and food and water ad libitum. Food and drink consumption was measured by highly sensitive sensors for automated online measurements located in the feeding and water ports. Energy expenditure (EE), O2 consumption and CO2 production, respiratory exchange rate (RER=VCO2/VO2) and locomotor activity are also measured. O2 and CO2 volumes are measured at the inlet ports of each cage through which a predetermined airflow is ventilated (0.4 L min^−1^) and compared regularly with a reference empty cage. EE was calculated using the Weir equation for respiratory gas exchange measurements. Ambulatory activity was measured with the help of an infrared light beam-based activity monitoring system. All sensors operate efficiently in both light and dark phases, allowing for continuous recording. Parameters were recorded every 5 min throughout the experiment.

Data analysis was performed using extracted raw values of VO2 consumed (expressed in mL h^−1^), VCO2 production (expressed in mL ^h−1^), energy expenditure (kcal/h), activity (cm), food intake (g) and drink (mL). The fatty acid oxidation has been calculated as followed (Bruss et al., 2010):

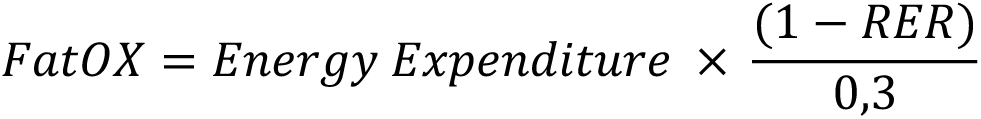

### Body composition analysis

Body mass composition was analyzed using an EchoMedical systems’ EchoMRI (RMN Analyzer minispeq mq7.5; Bruker Optik, Germany) in accordance with the manufacturer’s instructions. Readings of body composition were given within a 1-min time-frame. Measurements were performed 3 weeks after stereotactic delivery of the AAVs.

### Overnight fasting and re-feeding

Molecular and immunofluorescence analyses were assessed using mouse brains dissected following overnight fasting (18 H) and refeeding (2 H). The non-fasted group (NF) was killed at the same time point as the fasted group.

### Food intake measurement after either PTHrP or ghrelin injections

Mice were isolated in a regular housing cage for four days. Each day, food was removed at 9 am and mice were exposed to pre-weighted chow diet. After 2 H, the pellets were weighted and the mice received either subcutaneous injections of PTHrP (1 mg/kg) or saline solution, or intraperitoneal injections of ghrelin (1 mg/kg) or saline solution. 4 H after the injections, the pellets were weighted and the mice exposed to diet *ad libitum*. The two first days were considered as acclimatation days. The two last days were considered as testing days with saline injection the third day and PTHrP injection the fourth day.

### Food intake measurement after lalistat-1 and PTHrP intracerebroventricular injections

Mice were isolated in regular housing cage for three days. The first 2 days, mice received a first intracerebroventricular injection of saline solution at 6 pm and a second at 10 am and were exposed to pre-weighted chow pellets for 8 hours. The last days, mice randomly received DMSO (0,4%) or lalistat-1 (100 µg; 6098; Tocris) injection at 6 pm, and saline or PTHrP (40 ng; 4017147; Bachem) injection at 10 am and exposed to pre-weighted chow pellets for 8 hours.

### Food intake measurement after fasting

Mice were isolated in a regular housing cage for three days. The second day, mice were exposed to pre-weighted chow pellets for 30min at 10 am. At 6 pm, the food was removed for 16 H over-night fasting. The third day, mice were exposed to pre-weighted chow pellets for 30 min.

### Energy expenditure measurement

The experiment was performed 5 weeks after stereotaxic delivery of viral vectors and energy expenditure was measured using automated metabolic cages (Labmaster, TSE Systems GmbH, Bad Homburg, Germany). Mice were isolated in regular housing cages (35.5 x 18 x 12.5 cm) at 10 a.m.. They received an i.p. injection of saline solution at 7 p.m. for two consecutive days. The 3^d^ day at 10 a.m., mice were placed in automated metabolic cages and energy expenditure was measured for 48 H. During these 48 H of metabolic cage measurements, mice were injected (at 7 p.m.) with saline solution the 1^st^ day and PTH (1 mg/kg) the 2^nd^ day.

### Tissue collection

Mice allocated to immunofluorescence studies were deeply anesthetized with a mixture of ketamine hydrochloride (20mg/ml BW; 1000 Virbac) and xylazine (1000mg/ml BW; Rompun 2%; Bayer) and were perfused transcardially with a solution containing 0.9% NaCl, followed by 4% paraformaldehyde (PFA) in phosphate buffer (pH 7.3). Animals were sacrificed in randomized order to minimize experimental bias. The mice were then decapitated, and the brain was carefully removed. Mice allocated to molecular biology analysis were killed by cervical dislocation. The whole HpT was carefully dissected, immediately snap-frozen in liquid-nitrogen, and maintained at 80°C until further processing. The white adipose tissue (WAT) and brown adipose tissue (BAT) were harvested and were maintained in PFA 4% for 24 h before being sectioned and used for histological analysis.

### Slice Electrophysiology

Electrophysiological recordings were performed in non-fasted 6-9 week old AgRPCre:tdTomato mice. Mice were intracardially perfused under deep anesthesia (ketamine/xylazine, 100/10 mg/kg) with an ice-cold oxygenated (95% O2/5% CO2) perfusion solution that contained (in mM) 200 sucrose, 28 NaHCO3, 2.5 KCl, 7 MgCl2, 1.25 NaH2PO4, 0.5 CaCl2, 1 L-ascorbate, 3 Na-pyruvate and 8 D-glucose (pH 7.4). The brain was quickly removed and immersed in the same ice-cold oxygenated perfusion solution. Three to five 250 μm coronal slices containing the arcuate nucleus were obtained with a vibroslice (Leica VT1000S) and placed for 1 h at room temperature in an oxygenated recovery ACSF solution containing (in mM): 118 NaCl, 5 KCl, 1 MgCl2, 25 NaHCO3, 1.2 NaH2PO4, 1.5 CaCl2, 5 HEPES, 2.5 D-glucose and 15 sucrose (osmolarity adjusted to 310 mOsM with sucrose, pH 7.4). After recovery, slices were perfused with the same ACSF oxygenated media in a recording chamber placed under a microscope (Slicescope, Scientifica) outfitted for fluorescence and IR-DIC videomicroscopy. Viable arcuate AgRP neurons were visualized with a fluorescence video camera (Rolera Bolt Qimaging camera). For cell-attached recordings, borosilicate pipettes (4-6 MΩ; 1.5 mm OD, Sutter Instrument) were filled with filtered extracellular medium.

For the measurement of the firing rate of AgRP neurons in response to PTHrp (100nM), action potential frequency was quantified after a stable baseline was established before (control: over the last 120 to 60 sec before PTHrp application), during (stimulated by 100 nM PTHrp 5-7 min: over the last 60 sec of PTHrp application) and after (reversal 10 min: over 60 sec, 10 min after PTHrp application) PTHrp application, at room temperature. Recordings were made using a EPC10 Heka amplifier including data acquisition LIH 8+8 AD/DA interface and acquired at 2 kHz using patchmaster (2×91 (Harvard Bioscience, Inc., HEKA Elektronik GmbH). All acquired data were analyzed with Clampfit software (Molecular Devices, Inc.).

### Primary hypothalamic neuronal cultures

Hypothalamic cells were isolated from mouse embryos (embryonic day 16.5). After dissections, hypothalamus was digested with trypsin 0.05% and EDTA 0.02% for 15 min at 37°C. After three washes with DMEM (61965059; Thermo Fisher Scientific), supplemented 10% FBS, 100 U/ml penicillin-streptomycin, cells were dissociated and then plated. The dissociated cells were plated onto poly-L-lysine-coated plates or glass coverslips for microscopic examination. 24h after plating, the medium was replaced with Neurobasal medium (Thermo Fisher Scientific) containing B27 supplement (Thermo Fisher Scientific), GlutaMAX and Mycozap (Lonza), and was renewed two times per week while neurons were maintained in 5% CO2 and 37°C. To decrease the proportion of glial cells present in the culture, 2 µM of Ara-C inhibitor (C1768, Sigma) was added in pre-warmed complete Neurobasal medium, starting at Day In Vitro (DIV) 5. Experiments were performed at DIV12.

### N38 cell line culture

Murine hypothalamic neural (N38) cells (CLU118-P; Tebu-Bio) were grown in normal culture medium (DMEMC) comprising high glucose (25 mM) DMEM (D5796; Sigma), 10% FBS (30-2020; ATCC), 1% antibiotics (penicillin-streptomycin; 15140-122; Gibco) at 37°C and 5% CO_2_.

### Treatment of cell cultures

Primary hypothalamic cell treatments were administered in neurobasal medium. N38 cell treatments were administered in DMEMC. PTHrP (100 nM) and Bafilomycin A1 (1 mM) were administered for 4 hours. Lalistat-1 (10 µM) was administered 20 H before PTHrP treatment onset, for 4 hours. TCS21311 PKC inhibitor (75 nM) was administered 20 min before PTHrP treatment onset for 4 H. Ghrelin (500 nM) was administered for 1 hour.

### Quantitative RT-PCR

Tissues were immediately flash-frozen after dissection and total RNA was isolated with TRIzol Reagent using a homogenizer. Single-strand cDNA was synthesized from total RNA (2 μg) by using SuperScript II Reverse Transcriptase. qRT–PCR was performed using iTAQ SYBR Green (BioRad). Real-Time PCR System. Relative expression of *Pthrp* and *Pth1r* was calculated using the 2(11C.t/) method. Conditions for real-time PCR were: initial denaturation for 10 min at 95°C, followed by amplification cycles with 15 s at 95°C, and 1 min at 60°C. RT-PCR primers used were: *Pthrp* primer forward 5’– TGCCTCAGTCTTCTTGCCTTA–3’, *Pthrp* primer reverse 5’– CCCTAACCATGAACATTCTGT–3’, *Pth1r* primer forward 5’–CCTGAGTCAGCTCC TCAACC– 3’, *Pth1r* primer reverse: 5’–GCCGGCCCAATTGTGTGTC– 3’.

### Western blots

Brain tissue or hypothalamic cells were homogenized in RIPA lysis buffer (25mM Tris-HCl pH 7.6, 150 mM NaCl, 1% NP-40, 1% sodium deoxycholate, 0.1% SDS) (Pierce Thermo Scientific), supplemented with protease (cOmplete, Sigma) and phosphatase (phosSTOP, Sigma) inhibitors. Protein concentration was measured with Pierce BCA protein Assay Kit (ThermoFischer Scientific) prior to the Western blot assay. Tissue lysates were mixed with loading buffer and reducing agent (Invitrogen NP0004), and proteins were separated on 10%, 12,5% or 15% poly-acrylamide gel and subsequently transferred by semi-dry transfer onto a PVDF membrane (Trans-blot Turbo Mini PVDF, Biorad). The blots were blocked in 6% milk in Tris-buffered saline with Tween (TBS-T) or 2,5% Bovin Serum Albumin (BSA) and incubated with the primary antibodies, including rabbit polyclonal anti-AgRP (1/500, abcam, ab254558), mouse monoclonal anti-ATG5 (1/1000, Nanotools, 0262-100), mouse monoclonal anti-Beclin (1/1000, BD Transduction Laboratories, AB_399484), rabbit polyclonal anti-LC3 (1/5000, Sigma, L7543), guinea pig polyclonal anti-p62 (1/1000, Progen, GP62-C), rabbit polyclonal anti-PTH1r (1/1000, abcam, ab180762), mouse polyclonal anti-PTHrp (1/50, Santa-Cruz, sc-53936), mouse polyclonal anti-ATG5 (1:5000, Nanotools, 0262-100), rabbit polyclonal anti-VPS34 (1/1000, Cell Signaling, 4273), rabbit polyclonal pPKA (1/1000, CellSignaling, rabbit polyclonal 4781), PKA (1/1000, CellSignaling, 4782), rabbit polyclonal pPKCθ (1/1000, CellSignaling, 9377), rabbit polyclinal PKCθ (1/1000, CellSignaling, 13643) and mouse monoclonal anti-β-actin (1/5000, Sigma, A5316). To detect protein signal, the following Horseradish peroxidase–conjugated secondary antibodies were used: anti-rabbit IgG HRP-linked (1:5000, Cell Signaling, 7074), anti-mouse IgG HRP-linked (1/5000, Cell Signaling, 7076) and anti-guinea pig IgG HRP-linked antibody (1/5000, Sigma, A5545) revealed using an ECL kit (Clarity Western ECL Substrate: BioRad) for protein detection with a Chemidoc Imaging System (Biorad). Bands were quantified using the Image Lab software.

Homozygous AgRP^DTR/DTR^ mice (AgRP-KO) were euthanized by cervical dislocation and their heads were immediately cut and soaked in liquid nitrogen for 3-4 seconds. The brain was then removed and dissected on an ice-cold surface: the ARC was extracted to assess and compare the expression of AgRP, a 14kDa peptide synthesized by AgRP neurons, in the hypothalamus of AgRP-control vs. AgRP-ablated overnight fasted mice. ARC samples were sonicated and homogenized in 200 µl lysis buffer (ThermoFisher Scientific, Invitrogen, Tissue Extraction Reagent II – catalog #FNN0081) supplemented with protease inhibitor (ROCHE 05056489001, Inhibiteur de Proteases Complete, EDTA-free, diluted 250X). Brain samples were then centrifuged at 4°C for 20min. Aliquots (2µl) of the homogenates were added to 200µl of Bradford (Bio-Rad Protein Assay Dye Reagent Concentrate – cat. 5000006 – stored at 4°C) and were used for protein quantification. Equal amounts of proteins (30 µg/µl) for each sample were loaded onto 4-15% polyacrylamide gels (BIO-RAD mini-protean TGX, stain-Free Gels, cat #4568084; 4-15%). Proteins were separated by SDS-PAGE and transferred to PVDF membranes (Trans-blot Turbo Mini PVDF, #1704156, 0.2µm, Bio-Rad). Theses membranes were blocked in 5% milk for 1hr and were then incubated overnight with the following primary antibodies (diluted in 1% milk) at 4°C: rabbit anti-AgRP (1:1000, abcam, ab254558) and mouse monoclonal anti-β-actin (1:1000, Sigma, A5316). Membranes were then washed 3 times for 10 min in PBS-T (Tween 20, Sigma-Aldrich – P9416-100ml) and were incubated in HRP-coupled secondary antibody binding: anti-rabbit IgG HRP-linked antibody (1/5000, Cell Signaling, 7074) and anti-mouse IgG HRP-linked antibody (1/5000, Cell Signaling, 7076). Proteins were revealed and detected via enhanced chemiluminescence using an ECL kit (Clarity Western ECL Substrate – BIO-RAD - 1705061). Images were acquired with a GE Amersham Imager 600 (UV 29083463 Luminescent Image Analyzer).

### Hematoxylin and eosin staining and WAT adipocyte diameter analysis

Adipose tissue was fixed in 4% PFA then paraffin-embedded for H&E staining. Pictures of the sections were taken under light microscopy with a slide scanner (Nanozoomer 2.0 HT, Hamamatsu) using 20X objective. The images were then analyzed using the Adiposoft plug-in of Fiji software by counting at least 1,000 adipocytes per animals.

### Immunofluorescence

Perfused brains were cut at 40-micron thick coronal sections using a vibrating microtome (VT1000S, Leica). Immunostaining was performed on free-floating sections. Non-specific staining was blocked by 0.25% Triton and 10% donkey serum albumin (Sigma-Aldrich). Sections or cells were incubated with the following primary antibodies at 4°C: rabbit polyclonal anti-AgRP (1/200, abcam, ab254558), rabbit polyclonal aniti-LC3 (1/200, Sigma, L7543), guinea pig polyclonal anti-p62 (1/200, Progen, GP62-C), rabbit polyclonal anti-PTH1r (1/200, abcam, ab180762), mouse polyclonal anti-PTHrp (1/50, Santa-Cruz, sc-53936), MAP2 (1/200, Millipore, MAB364). Sections or cells were incubated with Alexa-conjugated secondary antibodiesè (1/800, ThermoFischer) for 2h at room temperature. Sections or cells were incubated with HCS LipidTox (1:200, ThermoFischer, H34476). Fluorescent sections or cells were stained with the nuclear dye Hoechst and then mounted using Fluoromount aqueous mounting medium (Sigma-Aldrich). Free-floating sections were washed 3 times for 10 min in PBS1X, incubated in 0.2% Triton X-100 solution for 20 min, rinsed again in PBS (3×10min), and blocked for 1hr in 3% BSA (Bovine Serum Albumin, Sigma-Aldrich, A2153). The slices were then incubated with the primary antibody anti-AgRP (1/200, abcam, ab254558) in a blocking solution of 1% BSA for 72hrs at 4 °C. They were later washed 3 times for 10 min in PBS1X before being incubated with the secondary antibody (diluted 1/500 in PBS1X) for 1hr at room temperature. Finally, the samples were rinsed 3 times for 10 min in PBS1X, stained with DAPI (diluted 1/10000 in PBS1X, Sigma D9542) for 10 to 15 min, washed twice for 10 min with PB1X (0.1M) and mounted onto microscope slides (SuperFrost^®^ Plus adhesive microscope slides, 25×75mm) with an antifade mounting medium (Vectashield^®^) before being cover slipped.

### *In situ* hybridization followed by immunofluorescence staining

*Pthrp* and *Pth1r* mRNA *in-situ* hybridization on mouse brain sections was performed with RNAscope Multiplex Fluorescent assay and ACD HybEZ^TM^ II Hybridization System (Biotechne). Mice were transcardially perfused with cold PBS, followed by 4% PFA as described above. Brains were post-fixed 24 H in 4% PFA, cryoprotected in 30% sucrose at 4°C, embedded in cryo-embedding medium (OCT), and frozen. 12 µm brain sections were processed according to the manufacturer’s instructions. RNAscope probes Mm-Pth1r (Cat N° 426191) and Mm-Pthrp-C2 (Cat. No. 456521-C2) were visualized with a confocal microscope (LSM 700, Zeiss) with a 20X immersion objective.

### Immunofluorescence imaging and quantification

Images of cells were acquired using an Apotome and images of tissues were acquired using a Spinning Disk, both with the Zen Imaging software (Zeiss). Z-stacks were obtained using sequential scanning. All analysis were performed using the Icy open-source platform (http://www.icy.bioimageanalysis.org). Cell counting values are expressed as the mean number of immune-positive cell counted. Fluorescence intensity values are expressed as the mean intensity of cell counted. Number of puncta are expressed as the mean of puncta of cell counted. The number of subjects, brain slices, primary neuron preparations included in each counting are mentioned in the corresponding legends. Confocal images (LSM 700, Zeiss) were obtained with a 10X immersion objective (Zeiss) and were visualized, using the ICY software for bioimage analysis (Image J), in order to evaluate AgRP expression in the ARC and PVN of AgRP-ablated vs. AgRP-control mice.

### Metabolic data analysis

For metabolomic analysis, the extraction solution was composed of 50% methanol, 30% acetonitrile (ACN) and 20% water. The volume of the extraction solution was adjusted to cell number (1 ml per 1E7 cells). After addition of extraction solution, samples were vortexed for 5 min at 4 °C and centrifuged at 16,000g for 15 min at 4 °C. The supernatants were collected and stored at −80 °C until analysis. LC/MS analyses were conducted on a QExactive Plus Orbitrap mass spectrometer equipped with an Ion Max source and a HESI II probe coupled to a Dionex UltiMate 3000 uHPLC system (Thermo). External mass calibration was performed using a standard calibration mixture every seven days, as recommended by the manufacturer. The 5 µl samples were injected onto a ZIC-pHILIC column (150 mm × 2.1 mm; i.d. 5 µm) with a guard column (20 mm × 2.1 mm; i.d. 5 µm) (Millipore) for LC separation. Buffer A was 20 mM ammonium carbonate, 0.1% ammonium hydroxide (pH 9.2), and buffer B was ACN. The chromatographic gradient was run at a flow rate of 0.200 µl min^−1^ as follows: 0– 20 min, linear gradient from 80% to 20% of buffer B; 20–20.5 min, linear gradient from 20% to 80% of buffer B; 20.5–28 min, 80% buffer B. The mass spectrometer was operated in full scan, polarity switching mode with the spray voltage set to 2.5 kV and the heated capillary held at 320 °C. The sheath gas flow was set to 20 units, the auxiliary gas flow to 5 units and the sweep gas flow to 0 units. The metabolites were detected across a mass range of 75–1,000 m/z at a resolution of 35,000 (at 200 m/z) with the automatic gain control target at 106 and the maximum injection time at 250 ms. Lock masses were used to ensure mass accuracy below 5 ppm. Data were acquired with Thermo Xcalibur software (Thermo). The peak areas of metabolites were determined using Thermo TraceFinder software (Thermo), identified by the exact mass of each singly charged ion and by the known retention time on the HPLC column.

Metabolomic data can contain a large portion of missing values, largely affecting the attribution of the metabolic features. A percentile transformation function (percentize in R) was used to normalize the data, assigning a rank to the value of the metabolite. In order to avoid direct assignment of 0 to highly variant data points, rank based normalization preserves the broad trends in the experiment compared to controls. For dimensional reduction, we performed a principal component analysis (PCA) on the normalized metabolite reads. We used the euclidean distance to determine orthogonal eigen vectors. prcomp function in R was used, and the first two principal components explaining the largest statistical variability in the data (approximately 70%) was plotted in a biplot representation with controls (Vehicle) and experimental samples (PTHrp). A heatmap representation of the metabolite was generated using the heatmap function in R. Furthermore, we clustered the data using a K-means clustering algorithm (function kmeans in R) to observe segregation of metabolites into distinct clusters. The number of k medians was set to 6, while the maximal iterations was at 10000. A “MacQueen” algorithm was used to cluster the data in order to reduce the sum of squared errors. Points are assigned to nearest centroid according to the Euclidean distance and the centroids are updated using the mean of the points in the group. The algorithm iterates till the minimum sum of squared errors is achieved.

## Statistical analysis

All data were expressed as mean ± SEM. Statistical analyses were performed using the Prism software (GraphPad, version 6), with p < 0.05 considered statistically significant. Data were analyzed using unpaired two-tailed Student’s *t*-test, unpaired one-tailed Mann-Whitney, unpaired Kruskall-Wallis followed by Dunn post-hoc test, one-way analysis of variance (ANOVA), or two-way ANOVA followed by Dunn post-hoc test.

## Supplementary Figure Legends

**Supplementary figure 1: *Pth1r* and *Pthrp* are abundantly expressed in the Arcuate nuclei (ARC) of the hypothalamus (HpT). A**) Representative co-immunofluorescence staining with PTH1R (orange) or PTHrP (red) antibodies performed on mouse coronal brain slices (40◻m) at the level of the HpT along the rostro-caudal axis, (from Bregma-1.5 to-5.0, with 0.7 interval between each representative sections). Representative brain coronal schemes are present on the top of each paired images. Mounting medium with *DAPI* was used for blue nuclear counterstaining.

**Supplementary Figure 2: *Pth1r* downregulation in POMC+ neuron has no incidence on energy metabolism. A**) Immunofluorescence staining of PTH1R (upper panel) and PTHrP (lower panel) in 3Mo *Pomc*^Cre:tdTomato^ mice. **B**) Representative images of AgRP immunostaining in the ARC and PVN of fasted AgRP-control and mice in which AgRP-neurons were ablated at neonatal stage (AgRP^DTR/+^) mice. **C**). Western Blot analysis of AgRP expression in the ARC of AgRP-control vs. AgRP KO (AgRP^DTR/DTR^) fasted mice **D**) Co-immunofluorescence staining was performed using PTH1R (red) and PTHrP (orange) antibodies on brain slices of mice injected with AAV-*Agrp*-shRNA-Scramble, AAV-*Agrp*-shRNA-*Pth1r* or AAV-*Agrp*-shRNA-*Pthrp.* Each AAV expressed eGFP (green) under the control of AgRP promotor. **E-F)** Bar graphs representing (overall mean and day-night mean) of food intake of AAV-*Agrp*-shRNA-*Pth1r*, AAV-*Agrp*-shRNA-*Pthrp* and Scramble mice, measured in automated metabolic cages over 5 days, following 2 days of habituation. These measurements show **G**) Bar plots representing energy expenditure (EE) and respiratory exchange ratio (RER) in AgRP-sh-*PTH1r* mice. Data are expressed as mean ± s.e.m. For body weight analysis and dark-light analysis of metabolic parameters, the two-way ANOVA was used followed by a Sidak post-hoc test. \**P* < 0.05, \*\**P* < 0.01, \*\*\**P* < 0.001.

**Supplementary Figure 3: Interactions between PTH1R and either PTH or PTHrP have distinct functions in the HpT control energy metabolism.**

**A)** Schematic representation of stereotactic injections in the ARC of AAVs carrying shRNA against Pthrp (driven by a U6 promoter), bar graphs representing the relative mRNA expression of mice stereotactically injected with AAV-U6-shRNA-Scramble (n=13) versus AAV-U6-shRNA-*Pthrp* (n=13) and representative images of mice downregulated for *Pthrp* in the ARC versus control. **B)** Body weight gain and epididymal fat pad of mice downregulated for Pthrp in the ARC versus control. **C)** Body composition measurements (by NMR analyzer) assessing relative body fat mass, lean mass and fluid volume of mice injected with either AAV-U6-shRNA-*Pthrp* (n=15) or AAV-U6-shRNA-Scramble (n=16). **D)** Bar-plot graphs representing (overall mean and day-night means) food intake and energy expenditure (heat) in U6-sh-Pthrp and Scramble mice, measured in the automated metabolic cages over 5 days, following 2 days of habituation. **E-F)** Bar plots representing water intake, respiratory exchange ratio (RER), locomotor activity (distance), oxygen consumption (VO2) and carbon dioxide production (VCO2) in mice that received PTHrP (40 ng/H) (n=12) (**E**) or PTH (40 ng/H) (n=12) (**F**) versus controls (NaCl (0,9%)) (n=9/12) via osmotic pumps (i.c.v., for 2 weeks before the tests). **G**) Firing rate in *AgRP*^Cre:tdTomato^ neurons in response to 100 nM PTHrP (n = 13 cells from 8 mice). Results are shown as mean ± s.e.m. For the electrophysiological analysis, one-way ANOVA of paired data, with Dunn’s multiple comparison test was performed. \**P* < 0.05, \*\**P* < 0.01, \*\*\**P* < 0.001.

**Supplementary Figure 4: PTHrP treatment in either primary HpT neurons or HpT cell line induces AgRP levels. A)** Representative Western Blot and quantification of AgRP levels performed in primary HpT neurons treated with PTHrP. β-actin was used as a loading control for each sample. **B-C)** Representative Western Blot and quantification of AgRP levels performed in mouse AgRP cell lines (N38) after treatment with either PTHrP (**B**) PTH (**C**) or vehicle. β-actin was used as a loading control for each sample. Data are expressed as mean ± s.e.m. The Mann–Whitney test was used. \**P* < 0.05, \*\**P* < 0.01, \*\*\**P* < 0.001.

**Supplementary Figure 5: PTHrP induces significant increase in the proportion of free fatty acids in primary hypothalamic neurons. A**) Heatmap representing free fatty acid relative abundance in vehicle-and PTHrP-treated primary HpT neurons. Analysis was performed using liquid chromatography-tandem mass spectrometry (LC-MS/MS). Representation of the metabolites was generated using the heatmap function in R. **B**) PCA biplot showing that the datasets collected after treatment with either vehicle or PTHrP are orthogonal to each other. **C)** K-means clustering methods among the experimental datasets yields 6 clusters of metabolites, with the sum of squares by cluster (between samples/total samples) explaining 96.5 % of the sum of errors. **D)** Clusters 1 and 6 represent most of the free fatty acids in the experimental samples. Fatty acids in the cluster 1 consist of Decanoic acid, Docosahexaenoic acid, Eicosapentaenoic acid, Oleic acid and Palmitic acid, while in cluster 6, the fatty acids are Linolenic acid, Linoleic acid, Myristic acid, Octanoic acid and Palmitoleic acid.

**Supplementary Figure 6: Induction of FFA availability by PTHrP is dependent on lipophagy in hypothalamic neurons. A**) Representative images of primary HpT cells infected with a lentivirus expressing the pH-sensitive tandem mRFP-GFP-LC3 reporter (fusion of mRFP, GFP and LC3) that allow the visualization of autophagosome (yellow) and autophagolysosome (red). Neurons were treated with PTHrP (100nM) or vehicle for 4 H. Autophagosome (mRFP+ / GFP+) and autophagolysosome (mRFP+ / GFP-) puncta were quantified on n=4 experimental replicates and on n ≥ 10 individual neurons per group for each analysis (each neuronal preparation derived from at least 12 embryonic HpT). Scale bar: 20 μm. **B**) Representative images of SQSTM1/p62 immunofluorescence staining (scale bars: 20 μm) and puncta quantification performed on primary HpT neurons transfected with a lenti-shRNA-*Pth1r* and treated with PTHrP (100nM) (n=3 independent neuronal cultures). Analysis was performed on n=5 experimental replicates; n ≥ 8 individual neurons per group used for each experimental replicate (each neuronal preparation was derived from at least 12 embryonic HpT). Results are given as mean ± s.e.m. One-way ANOVA followed by Dunn’s post-hoc and Mann–Whitney tests were used for statistical analysis. \**P* < 0.05, \*\**P* < 0.01, \*\*\**P* < 0.001. NS: non-significant.

**Figure 7: PT**HrP **is involved in the transmission of feeding behavior induced by ghrelin. A**) Representative western blot and quantification of LC3 in the hypothalamus after ghrelin (3 mg/kg, n=5) or vehicle (n=8, NaCl 0,9%) i.p. injection. **B**) Representative western blot and quantification of p62, LC3 and PTHrP in N38 cells treated with ghrelin (500 nM for 1 H, n=5) or vehicle (NaCl 0,9%, n=5). Data are expressed as mean ± s.e.m. For western blot and PTHrP counting analysis, Mann-Whitney test was used. For LipidTox puncta, AgRP relative intensity and food intake measurement analysis, one-way ANOVA followed by Tuckey’s post-hoc test was used. *P < 0.05, **P < 0.01, ***P < 0.001.

