## Supplementary figures and images for "Parathyroid Hormone Receptor 1 Signaling Encodes Multifaceted Metabolic Programs in AgRP Neurons"

### Supplemental Figures

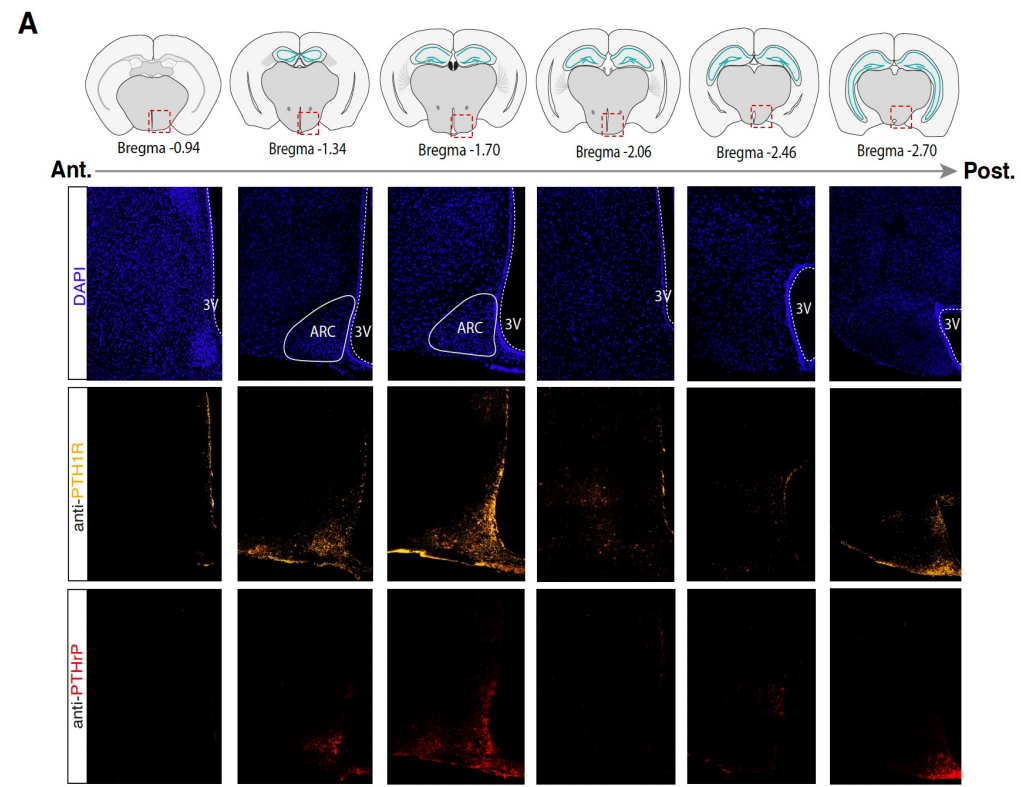

Figure S1

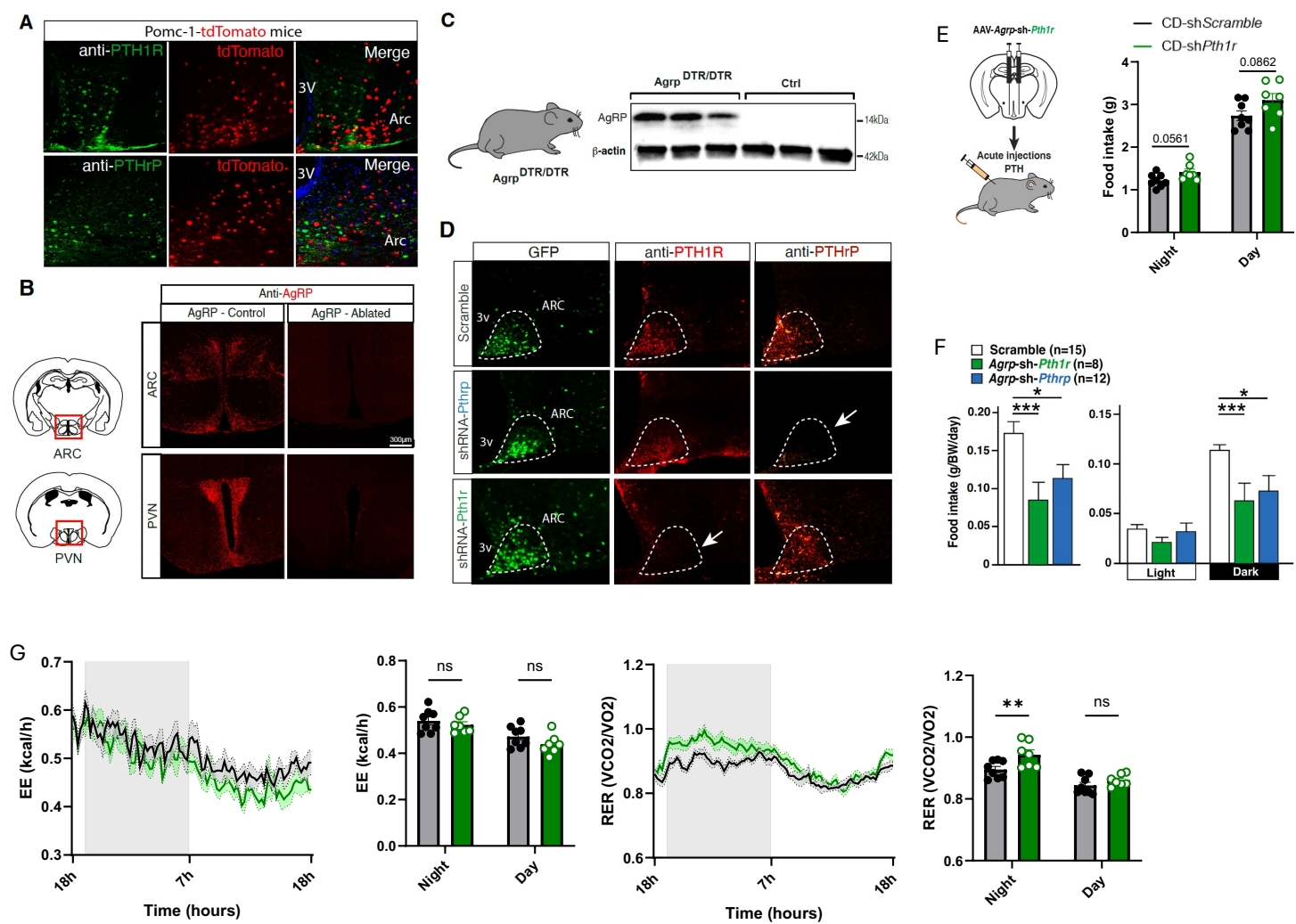

Figure S2

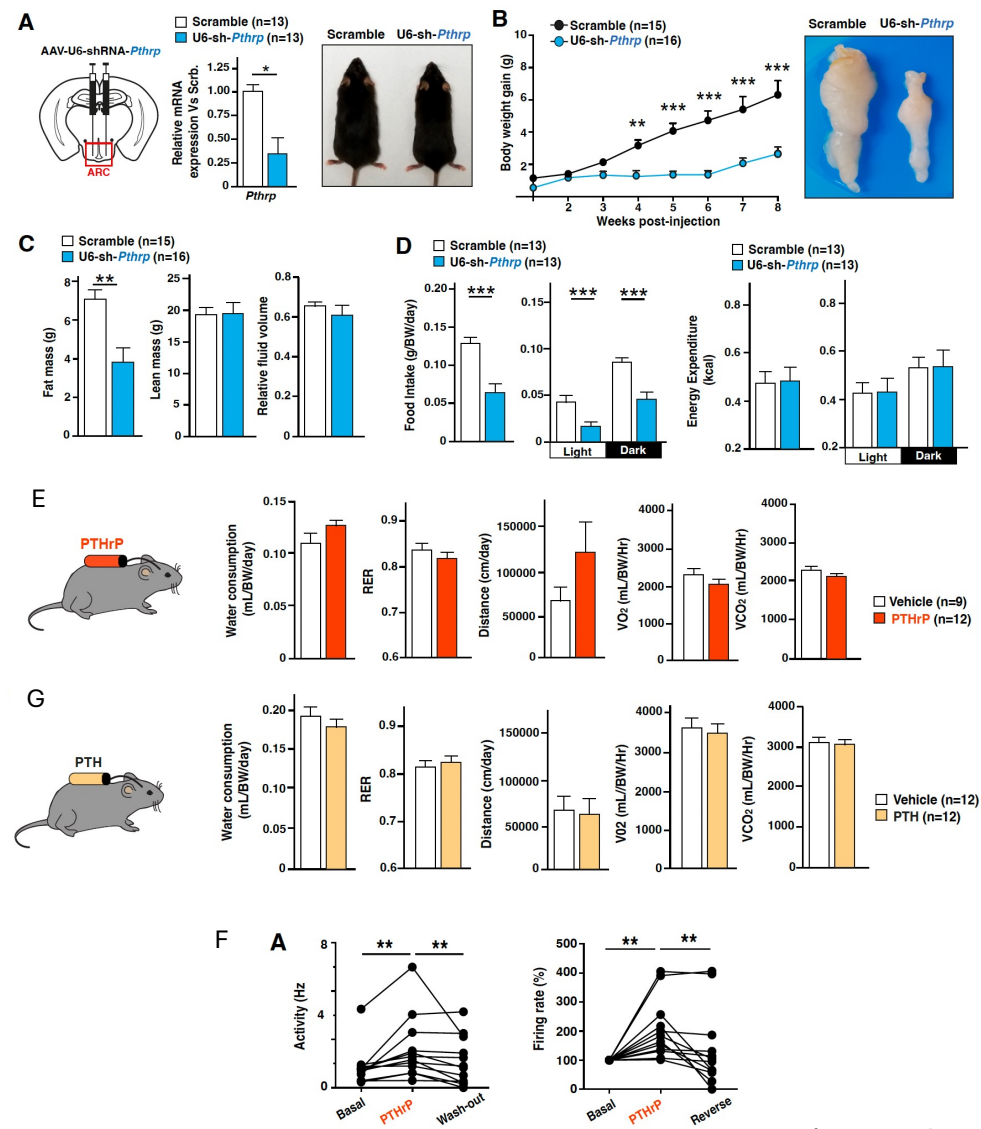

Figure S3

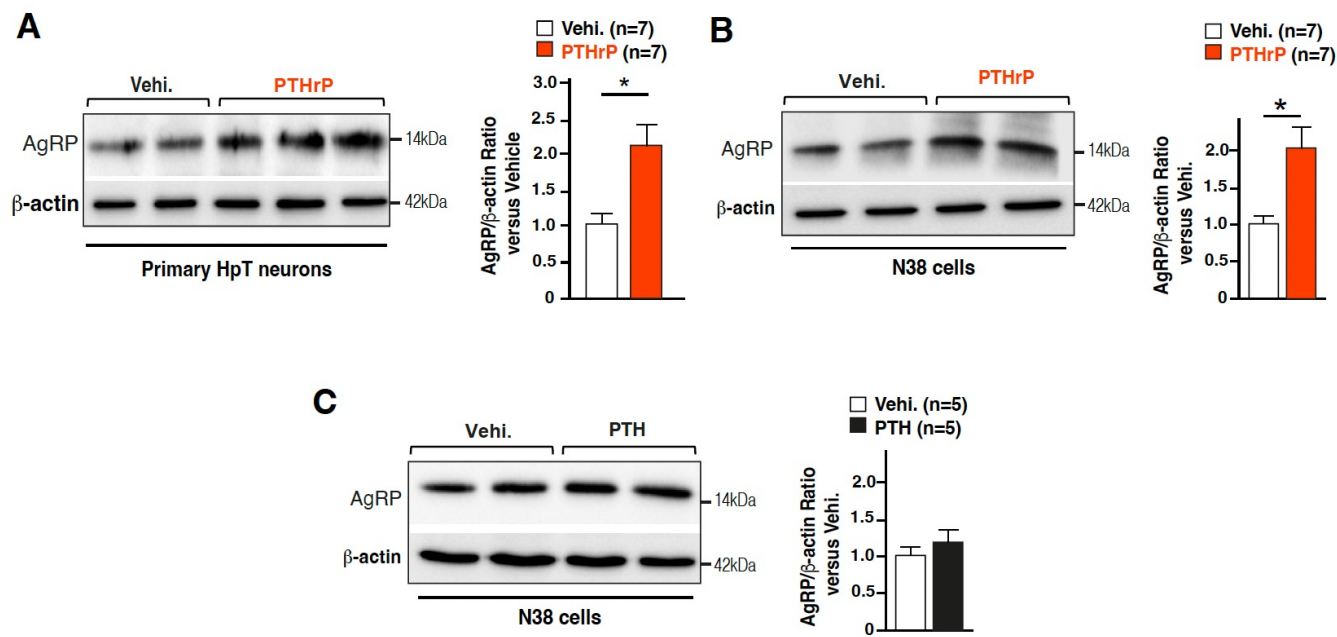

Figure S4

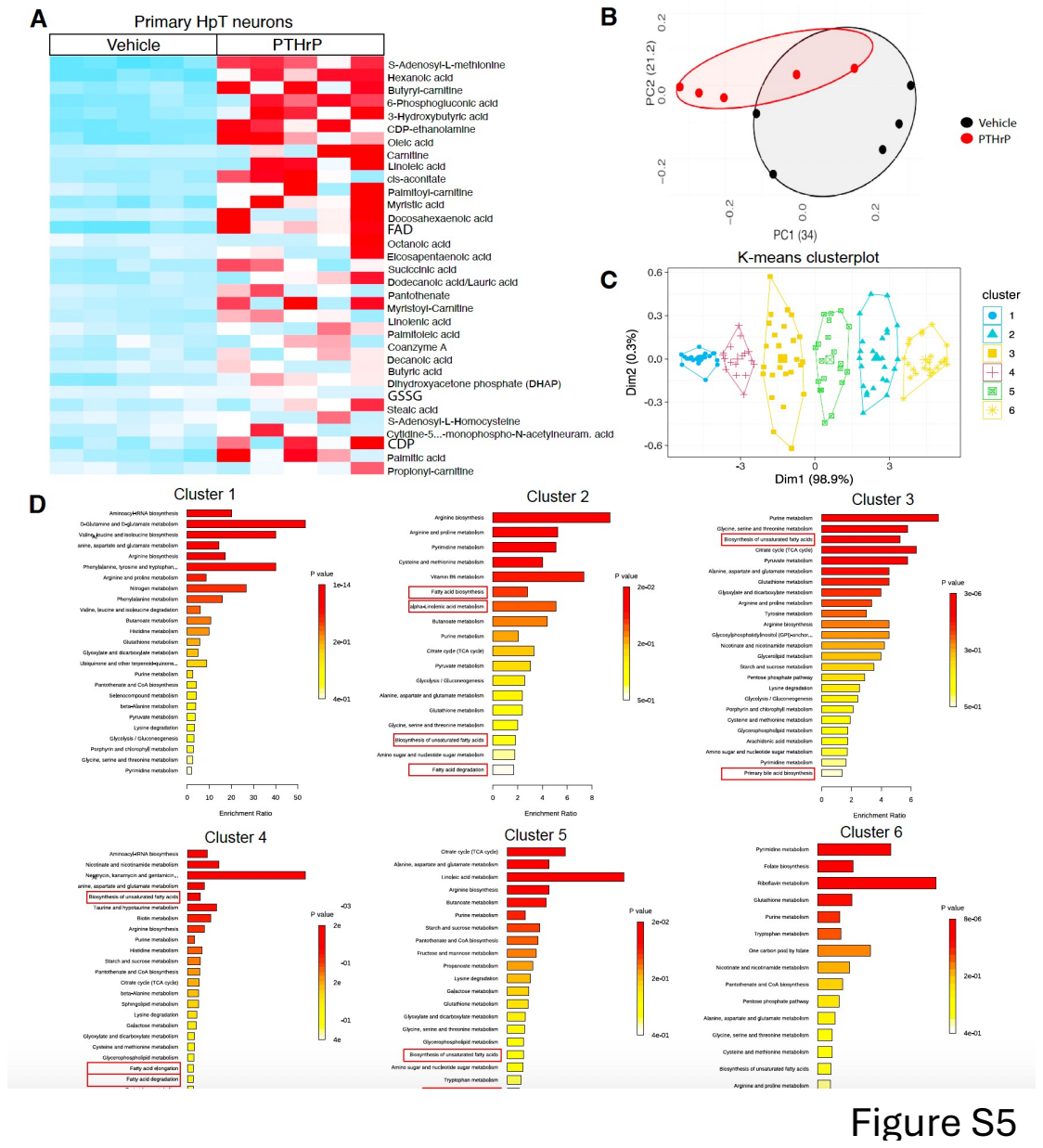

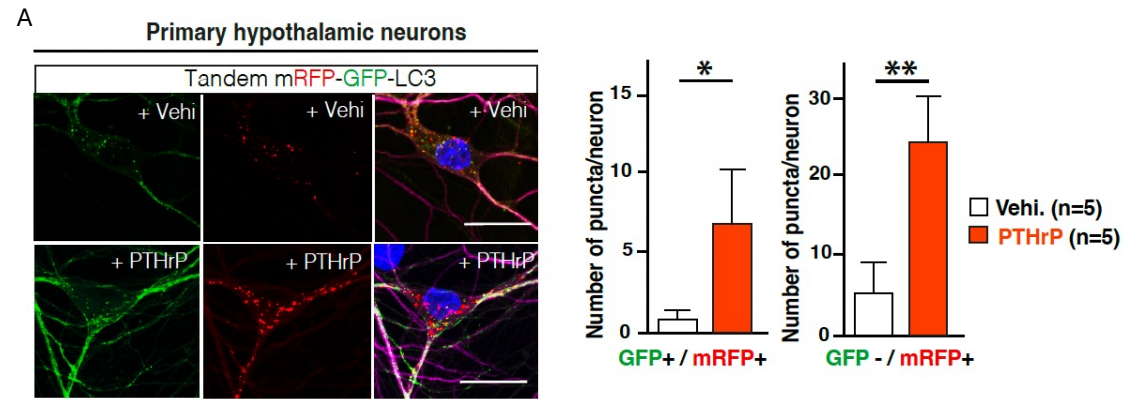

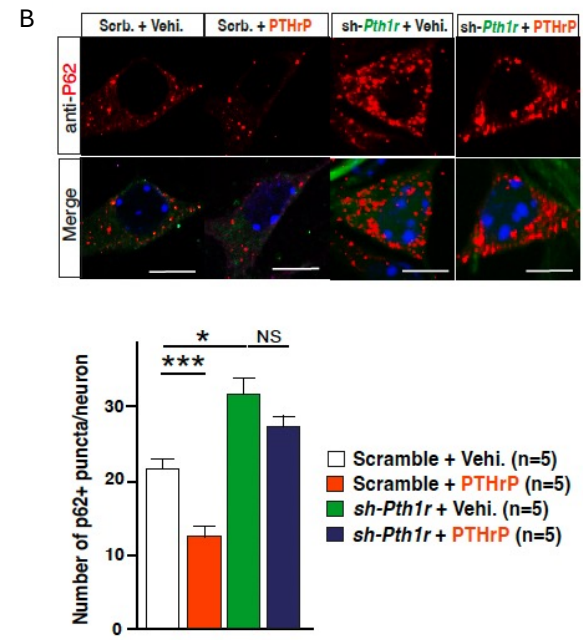

Figure S6

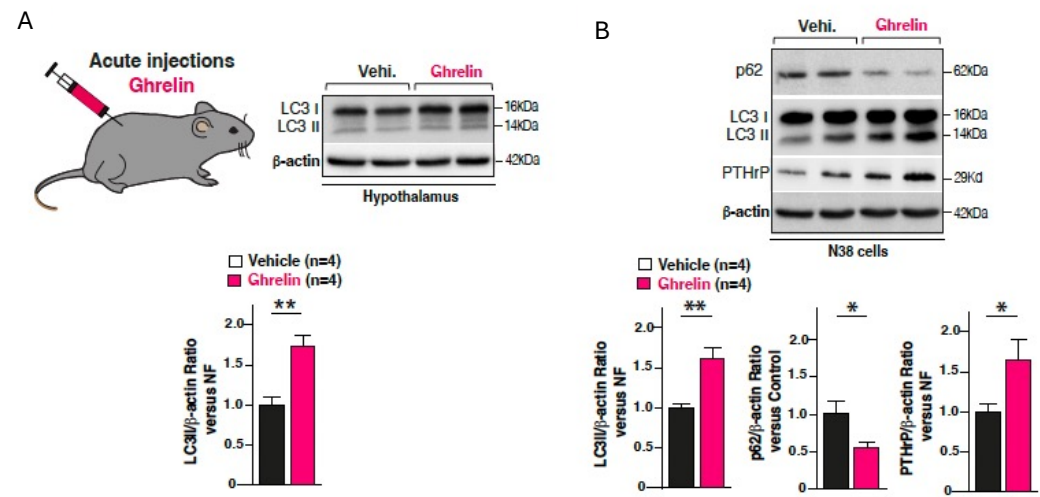

Figure S7
